# RBProximity-CLIP Enables Subcellular Mapping of RNA–Binding Protein Interactions at Nucleotide Resolution

**DOI:** 10.64898/2025.12.12.693770

**Authors:** Iwona Nowak, Ahsan H. Polash, Hang T. Huynh, Mahekdeep Kaur, Vivian Lobo, Jérémy Scutenaire, Michelle Fong, Ghaliah Alluhaibi, Dimitrios G. Anastasakis, Markus Hafner, Daniel Benhalevy, Aishe A. Sarshad

**Author notes:** Equal first author contribution (Iwona Nowak, Ahsan H. Polash, Hang T. Huynh).

## Abstract

RNA-binding proteins (RBPs) enable post-transcriptional gene regulation (PTGR) through specific interactions with RNA molecules, influencing processes ranging from nuclear processing and export to cytoplasmic localization, translation, storage and degradation. A key determinant of PTGR processes is the subcellular compartmentalization of RBPs, which dictates RNA targets they can access and the regulation performed in that environment. To characterize RBP-RNA interactions at subcellular resolution, we developed RBProximity-CLIP. RBProximity-CLIP enables compartment-specific isolation and profiling of individual RBP-RNA interactions by combining APEX2-based proximity labeling and 4-thiouridine–enhanced RNA-protein crosslinking, with sequential RBP- and biotin-affinity purifications. Using this approach, we profiled the RNA targets of three RBPs, AGO2, YBX1, and ELAVL1, across the cytoplasmic, nuclear, and nucleolar compartments, revealing nucleus-specific miRNA-mediated AGO2 targets, as well as subsets of YBX1 and ELAVL1 targets that differ by compartment, yet share identical binding motifs. RBProximity-CLIP enables specific and sensitive detection of compartment-specific RBP-RNA interactomes, thereby providing new insight into spatial gene regulation by RBPs.

## Background

Protein-RNA interactions enable post-transcriptional gene regulation (PTGR), affecting RNA transcription, processing, translation, stability, and subcellular localization [1]. RBPs typically recognize RNA through a limited set of RNA-binding domains (RBDs) that bind to short, but specific sequence or structural motifs on target RNAs [2, 3]. While RBPs evolved to molecularly recognize specific RNA motifs [4], the functional output of the RBP-RNA interaction often depends on the RBP’s subcellular localization, which also governs their access to particular RNA targets [5–11]. Given the importance of protein-RNA interactions for cellular homeostasis and disease [1, 5, 12], RBP expression, subcellular localization, and RNA binding are tightly regulated.

To date, protein-RNA interactions have primarily been studied by combining protein-RNA crosslinking with immunoprecipitation (CLIP) of the RBP, followed by RNA sequencing. All CLIP-based methods have their unique advantages and drawbacks [13]. In photoactivatable ribonucleoside-enhanced-CLIP (PAR-CLIP), crosslinking efficiency can be boosted by labeling nascent RNAs with the photoreactive nucleoside analogue 4-thiouridine (4SU), which can be crosslinked with amino acid side chains from interacting proteins using long-wavelength ultraviolet (UV) light [14]. Only crosslinked 4SU further results in a structural change that is misread during reverse transcription, leading to cDNA libraries containing T-to-C conversions at authentic crosslinking sites. This simplifies data analysis and enables the computational removal of abundant, non-crosslinked background derived from abundant RNA fragments [14]. Nevertheless, all standard CLIP techniques share a common limitation: RBPs are immunoprecipitated from whole-cell extracts, eliminating insight into the cellular location where the protein-RNA interactions occurred.

To increase their subcellular resolution, RIP- and CLIP-based techniques have successfully been combined with biochemical fractionation [6, 7, 15–17], which has enabled the study of subcellular RBP interactions [18]. Despite the development of protocols for purifying various subcellular compartments, these methods are limited to fractionable compartments and often suffer low signal-to-noise ratio [18, 19]. In addition, biochemical fractionation remains challenging, time-consuming, and highly user-dependent, resulting in technical variability even for established, fractionable compartments such as the cytosol, endoplasmic reticulum, and mitochondria [18, 20, 21].

Proximity-labeling and proximity-ligation methods overcome many challenges of biochemical fractionation and can be used for studying the subcellular localization of proteins and RNAs [10, 12, 22–32]. APEX2, an engineered soybean ascorbate peroxidase 2 [26], catalyzes the oxidation of biotin-phenol to generate spatially-restricted biotinylation, enabling rapid proximity-labelling [26, 28, 33]. The short-lived reactive species (< 1 ms) restrict biotinylation to the immediate vicinity of APEX2, within a reaction time of 30-60 seconds, ideal for dynamic processes [12, 26, 34]. APEX2 can be fused to specific localization elements, enabling compartment-specific biotinylation [35, 36]. This allows for proteomic and transcriptomic studies across a wide range of subcellular compartments, including those inaccessible to biochemical fractionation, e.g., the nucleolus, nuclear lamina, nuclear pore, cell:cell interface, and cell membranes [10, 26, 28, 29, 37, 38]. This versatility enabled diverse applications, including APEX-MS [34], APEX-RIP [30], APEX-seq [28, 29], and Proximity-CLIP [10].

By combining APEX2-mediated proximity labelling with UV-mediated protein–RNA crosslinking, and then footprinting the bound RNA elements, Proximity-CLIP profiles both the localized transcriptome and the specific RBP-bound RNA elements within any given subcellular compartment [10, 39, 40]. However, it does not enable characterization of RBP-specific subcellular targets, which to date has only been shown using Colocalization-CLIP [41]. Here, we present the development of RBProximity-CLIP, a method that enables the capture and characterization of RNA elements bound by individual RBPs at subcellular resolution. In RBProximity-CLIP, cellular compartments (e.g., cytoplasm, nucleus, or nucleoli) are biotinylated as in Proximity-CLIP using localized APEX2 elements. However, sequential affinity purification steps enable the isolation of RBP-specific interactions from a defined cellular compartment. We apply RBProximity-CLIP to dissect the subcellular RNA targets of three RBPs, Argonaute 2 (AGO2), Y-box binding protein 1 (YBX1), and Human antigen R (ELAVL1). RBProximity-CLIP approach enables precise RNP detection and size selection, resulting in enhanced sensitivity, specificity, and greater versatility compared to other approaches.

## Results

### Targeting cytoplasmic and nuclear AGO2–RNA interactions in HEK293 cells

We developed RBProximity-CLIP to expand the utility of CLIP methods, incorporating spatial resolution to enable the identification of RNA-RBP interactions in specific cellular microenvironments and to reveal context-dependent RBP functions. RBProximity-CLIP combines APEX2-mediated biotinylation and UVA/UVB-mediated protein-RNA crosslinking, followed by sequential biotin- and RBP-affinity purifications. Then RNA elements bound by compartment- and RBP- specific ribonucleoproteins (RNP) are analyzed as in fPAR-CLIP [42] (**Fig. 1A**). The RBProximity-CLIP workflow includes: 1) proximity biotinylation in live cells, 2) UV-induced protein-RNA crosslinking, followed by 3) immunoprecipitation (IP) to immobilize the RBP of interest, 4) RNase digestion of unprotected RNA and 3’ adaptor ligation to the remaining RNA footprint, 5) elution of the immobilized RNP and SDS-PAGE separation; 6) guided by the 3’-adapter fluorescence, the RBP-RNP is excised from the gel, 7) reconstituted into a soluble form by buffer exchange, followed by 8) capture of biotinylated RNPs by streptavidin affinity purification, and 9) release of compartment- and RBP-specific crosslinked RNA footprints by Proteinase K digestion, which are 10) transformed into a small RNA cDNA library for NGS analysis.

**Figure 1.**
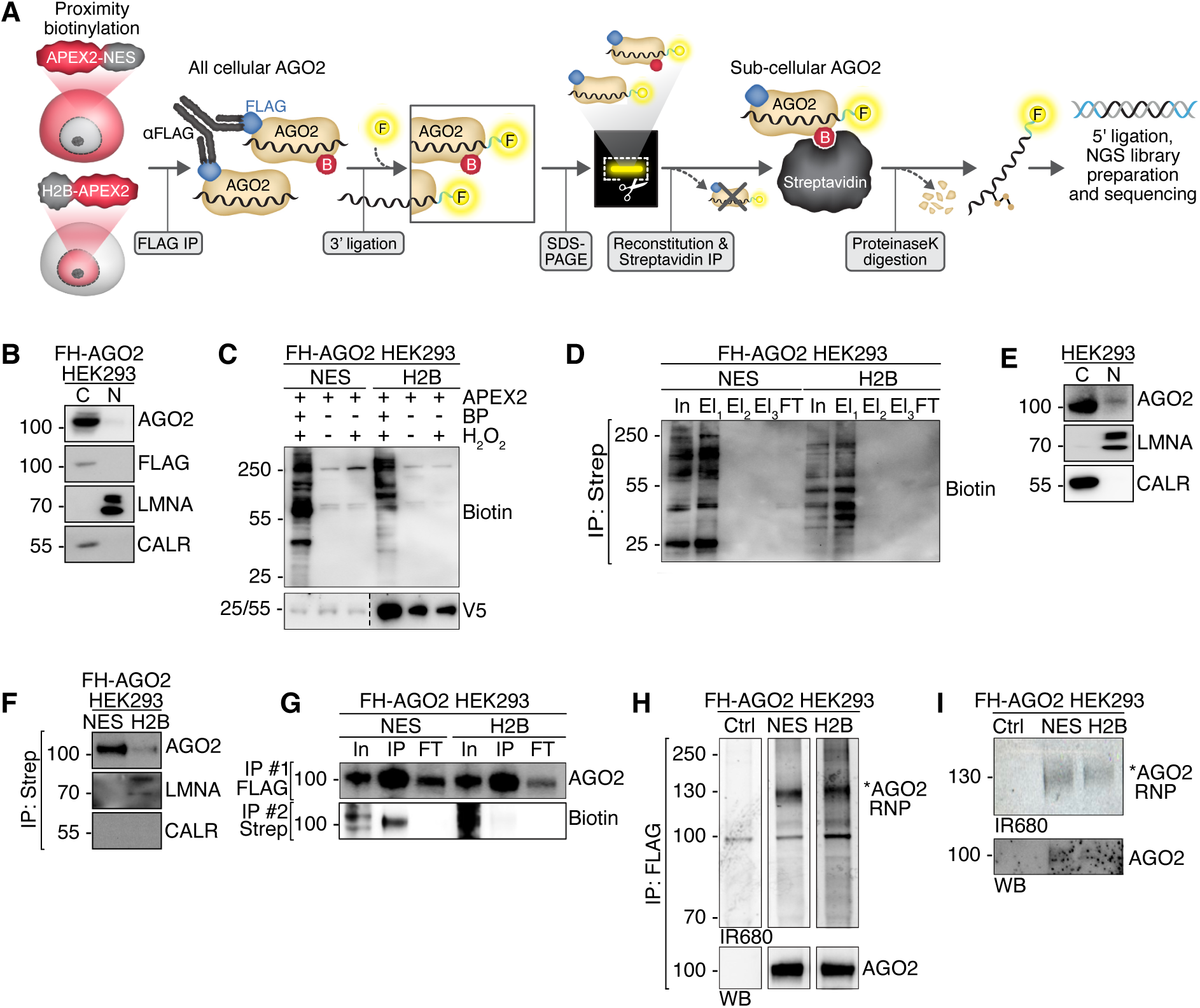
Cytoplasmic and nuclear RNA interactome of AGO2 in HEK293 cells. **(A)** Schematic overview of the RBProximity-CLIP workflow. HEK293 cells expressing APEX2–NES or APEX2–H2B undergo proximity biotinylation to label proteins in the cytoplasmic or nuclear compartment, respectively. FH-AGO2 is first isolated by anti-FLAG immunoprecipitation (IP), followed by 3′ fluorescent adapter ligation and SDS–PAGE separation. AGO2–RNA complexes are fluorescently visualized, gel-excised, reconstituted into solution and subjected to Streptavidin affinity purification for capture of biotinylated, compartment-specific AGO2 complexes. Then, AGO2-bound RNAs are recovered by proteinase K digestion, ligated to a 5′ adapter, and further processed into next-generation sequencing (NGS) libraries. **(B)** AGO2, FLAG, Lamin A/C (LMNA) and Calreticulin (CALR) Western blot analysis of biochemically fractioned cytoplasmic (C) and nuclear (N) preparations from HEK293 cells stably expressing FH-AGO2. **(C)** Biotin and V5 Western blot analysis of FH-AGO2 expressing HEK293 whole cell lysates in the presence or absence of proximity biotinylation components: V5-APEX2–NES (NES) or V5-APEX2–H2B (H2B), doxycycline-induced expression, biotin-phenol (BP) and hydrogen peroxide (H_2_O_2_). V5-APEX2–NES and V5-APEX2–H2B migrate at approximately 25 kDa and 55 kDa, respectively. **(D)** Biotin Western blot of Streptavidin affinity capture. Inputs comprise extracts of HEK293 cells stably expressing FH-AGO2 and either V5-APEX2–NES (NES) or V5-APEX2–H2B (H2B), after proximity biotinylation. Input (In), sequential elutions (El1, El2, El3) and flowthrough (FT). **(E)** AGO2, Lamin A/C (LMNA) and Calreticulin (CALR) Western blot analysis of biochemically fractioned cytoplasmic (C) and nuclear (N) preparations from HEK293 cells. **(F)** AGO2, Lamin A/C (LMNA) and Calreticulin (CALR) Western blot analysis of the material eluted after Streptavidin affinity capture from extracts of HEK293 cells stably expressing FH-AGO2 and either V5-APEX2–NES (NES) or V5-APEX2–H2B (H2B), after proximity biotinylation. **(G)** AGO2 and Biotin Western blot analysis of sequential FLAG IP (FH-AGO2) and Streptavidin affinity capture from HEK293 cells stably expressing V5-APEX2–NES (NES) or V5-APEX2–H2B (H2B), after proximity biotinylation. **(H)** Fluorescence imaging after SDS-PAGE of 3′-ligated AGO2-RNA complexes IP preparations (top) and a corresponding AGO2 Western blot analysis (bottom). Proximity biotinylation and FLAG-AGO2 IP were performed for HEK293 cells stably expressing FH-AGO2 and either V5-APEX2–NES (NES) or V5-APEX2–H2B (H2B). **(I)** Fluorescence imaging (top) and AGO2 Western blot analysis (bottom) of 3′ adapter–labeled AGO2–RNPs corresponding to 2.5% of the material eluted from the gel shown in (**H**).

We first applied RBProximity-CLIP to isolate nuclear and cytoplasmic targets of the main component of the RNA-induced silencing complex (RISC), AGO2. AGO proteins use small RNAs (miRNAs, siRNAs) to guide the RNA-induced silencing complex to partially complementary target RNAs [43]. The analysis of RNAs bound by the AGOs has been the subject of many studies [43]. In mammalian cells, including HEK293 cells, AGO proteins localize to the cytoplasm; nevertheless, in stem cells, germ cells, and in certain cancers, the AGOs can be found in considerable amounts in the nucleus, where they can engage in target RNA regulation using their effector machinery [6, 7, 15, 44–47].

We chose to first benchmark our method by profiling AGO2 binding to target RNAs across distinct subcellular compartments in HEK293 cells, expecting to capture robust miRNA-target sites mainly in the cytoplasm, where AGO2 is primarily localized (**Fig. 1B**). We developed stable HEK293 cells expressing FLAG-HA-tagged AGO2 (FH-AGO2) together with doxycycline-inducible APEX2 targeted either to the nucleus by fusion to Histone H2B (APEX2-H2B) or cytoplasm by fusion to a nuclear export signal (APEX2-NES) [10] (**Fig. 1A**). We validated APEX2-dependent biotinylation in each of the cell lines (**Fig. 1C**). A smear corresponding to biotinylated proteins was detected only in samples treated with the APEX2 substrates (biotin phenol- and H_2_O_2_) (**Fig. 1C**). Biotinylated proteins could be effectively captured by streptavidin affinity purification (**Fig. 1D**).

To compare the nucleocytoplasmic resolution achieved by proximity biotinylation versus biochemical fractionation, we examined the distribution of AGO2 relative to nuclear Lamin A/C and Calreticulin (an Endoplasmic Reticulum (ER) protein) in samples obtained by these two methods (**Fig 1E,F**). Both techniques showed exclusive detection of AGO2 in the expected cytoplasmic fractions, along with nuclear detection of Lamin A/C (**Fig 1E,F**). As expected, for APEX2-mediated cytoplasmic biotinylation, no biotinylated ER-lumen Calreticulin was detected (**Fig 1F**), in contrast to biochemical fraction (**Fig 1E**). We then performed FLAG-IP followed by sequential streptavidin affinity capture, confirming that this approach can selectively isolate cytoplasmic AGO2 (**Fig. 1G**).

After validation of the technical setup, we proceeded with performing the RBProximity-CLIP experiment. Cells underwent proximity-biotinylation and protein-RNA crosslinking, and FH-AGO2-RNPs were immunoprecipitated, treated with RNase T1, followed by ligation of a fluorescent adapter oligonucleotide to the 3’ends of protected RBP-bound RNA fragments. The ligated material was eluted, separated by SDS-PAGE, and visualized by fluorescent imaging, which confirmed successful AGO2-RNP capture migrating at **∼**130 kDa (**Fig. 1H, top**), whereas RNA-free AGO2 migrated at **∼**100 kDa (**Fig. 1H, bottom**). Subsequently, AGO2-RNPs were excised from the gel, reconstituted into solution (**Fig. 1I**), and subjected to streptavidin affinity purification for selective capture of compartment-specific AGO2. Nuclear and cytoplasmic AGO2-bound RNAs were eluted by Proteinase K digestion and processed into small RNA cDNA libraries. Consistent with the absence of AGO2 from the nucleus of HEK293 cells (**Fig. 1E,F**), we recovered only low amounts of cDNA libraries from nucleus-captured AGO2-RNPs, supporting the spatial specificity of RBProximity-CLIP.

### RBProximity-CLIP for AGO2 captures *bona fide* miRNA binding sites

RBProximity-CLIP libraries were sequenced, analyzed, and compared to AGO2 Fractionation-CLIPs from cytoplasmic and nuclear fractions of HEK293 cells [7]. Biological replicates of all conditions indicated reproducibility (⍴ = 0.56, 0.16, 0.69, 0.41, 0.72) (**Fig. 2A)**. Thus, for the following analyses, we merged the data from biological replicates and used the combined datasets to define AGO2 binding sites (clusters) based on overlapping sequence reads containing T-to-C mutations induced by UV-crosslinking of 4SU-incorporated RNA [14]. A cross-sample correlation analysis showed that cytoplasmic and nuclear profiles were more similar in the biochemically fractionated samples than in the RBProximity-CLIP samples, indicating that RBProximity-CLIP achieves a clearer separation between compartments (**Fig. 2B**). With RBProximity-CLIP, 5,712 and 585 clusters were identified for cytoplasmic and nuclear AGO2, respectively, reflecting roughly a 10-fold increased detection of cytoplasmic over nuclear signals (**Fig. 2C**). In comparison, the Fractionation-CLIP approach yielded 6,459 clusters in the cytoplasmic fraction and 1,887 in the nuclear fraction, presenting a more even detection in the two compartments (3.4-fold difference) (**Fig. 2C**), which aligns with experimental noise of biochemical fractionation, as previously reported [18, 19, 48]. Sensitivity for RBProximity-CLIP and the fractionated samples were overall lower than in whole-cell AGO2 fPAR-CLIP (**Fig. 2C**), implying a sensitivity cost for spatial specificity.

**Figure 2.**
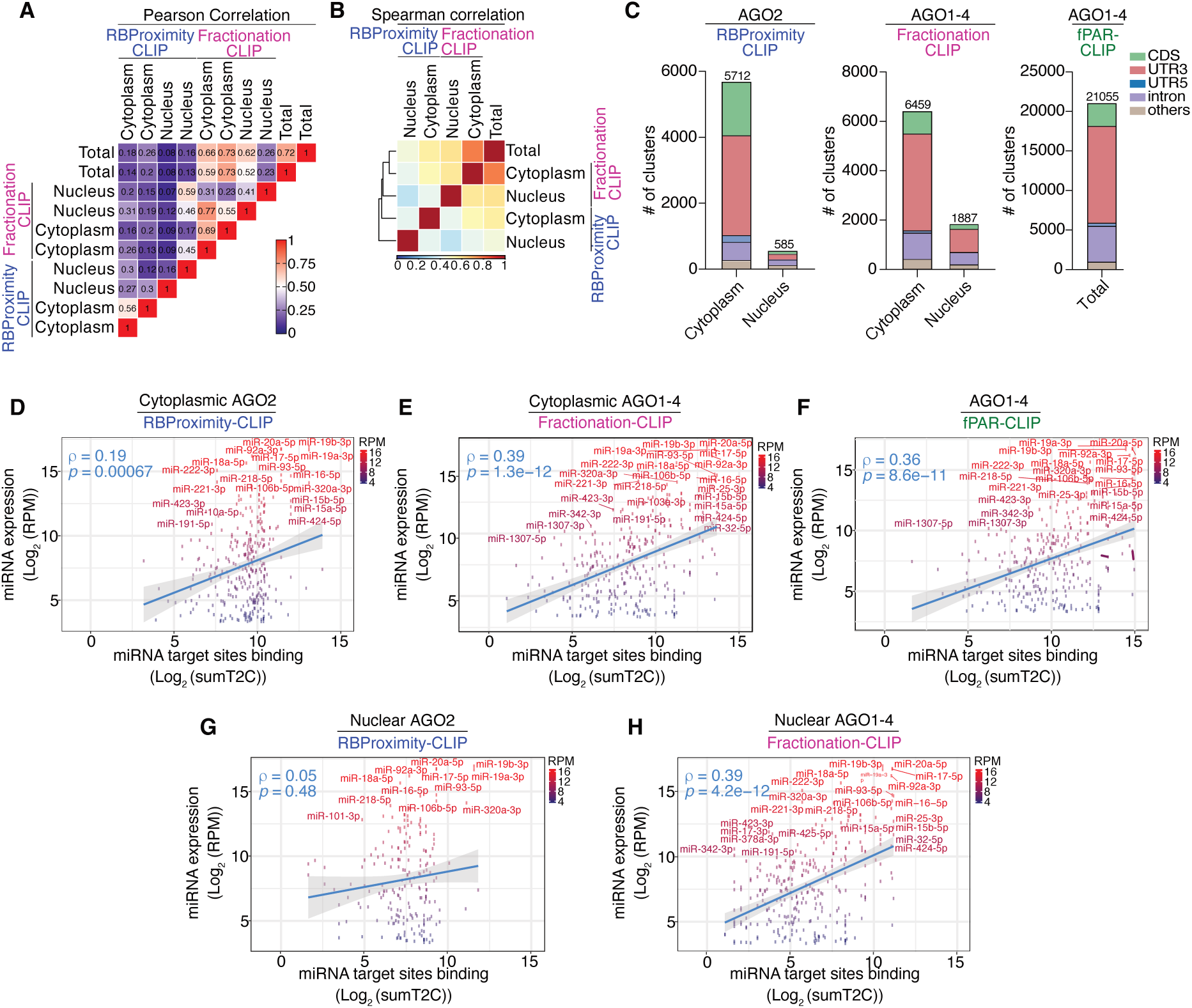
Comparative assessment of AGO2–RNA Interactions Across Subcellular Compartments Revealed by RBProximity-CLIP and Fractionation-CLIP. (A) Pearson correlation heatmap of transcript abundance across RBProximity-Fractionation-and whole-cell fPAR-CLIP datasets. **(B)** Spearman correlation heatmap of transcript abundance across RBProximity-CLIP Fractionation- and whole-cell fPAR-CLIP datasets, after pooling of replicates. Hierarchical clustering demonstrates the relationship between sample types. **(C)** Distribution of AGO2 binding sites (clusters) between transcriptome annotations, obtained by RBProximity- Fractionation- and whole-cell fPAR-CLIP. Coding sequences (CDS), 3′ untranslated regions (UTR3), 5′ untranslated regions (UTR5), introns, and all other annotations. **(D-F)** Correlation between number of crosslinked reads mapping to miRNA target sites, obtained by cytoplasmic (V5-APEX2–NES) RBProximity-CLIP (**D**), cytoplasmic Fractionation-CLIP (**E**) or whole-cell fPAR-CLIP (**F**), and miRNA expression obtained by miRNA-Seq. The blue line represents the linear regression fit with 95% confidence interval. The Spearman correlation coefficient (ρ) and corresponding *p*-value are indicated. Highly abundant miRNAs are labeled. # of reads with T to C conversion (sumT2C); Read per million (RPM). **(G,H)** Same as in (**D,E**), just for nuclear (V5-APEX2–H2B) RBProximity-CLIP (**G**) and nuclear Fractionation-CLIP (**H**) datasets.

Cytoplasmic RBProximity-CLIP and Fractionation-CLIP both showed a clear enrichment of AGO2 binding sites along mRNA 3’untranslated regions (UTR) (**Fig. 2C**), where most functional miRNA target sites are known to reside [43], suggesting that both approaches captured functional AGO2-RNA interactions. Nuclear RBProximity-CLIP profile differed markedly from the cytoplasmic fraction, with nearly tenfold fewer captured clusters with RBProximity-CLIP, consistent with the very low protein level of AGO2 in HEK293 cells nuclei (**Fig. 1E,F**). The nuclear fraction of Fractionation-CLIP showed a higher number of clusters and pronounced 3’ UTR binding, likely reflecting cytoplasmic leakage during the fractionation procedure (**Fig. 2C**), underscoring an advantage of fractionation-independent approaches such as RBProximity-CLIP that minimize sample manipulation and thus technical noise.

miRNA target sites are complementary to the miRNA ‘seed’ region (miRNA positions 2-7 or 2-8) [43]. To test whether spatial specificity maintains authentic detection of mRNA loading with the RISC complex, we tested whether miRNA expression levels correlate with occupancy of seed-sequence complementary elements along AGO2 binding sites obtained by fPAR-CLIP and RBProximity-CLIP. Our results confirm that the most expressed miRNAs are complementary to detected AGO2 binding sites obtained by fPAR-CLIP and RBProximity-CLIP, which are also enriched with T-to-C conversions that indicate authentic crosslinking events (**Fig. 2D-F**). Several highly expressed miRNAs, including hsa-miR-20a-5p, hsa-miR-19a-3p, and hsa-miR-16-5p, all from the miR-17∼92 cluster, displayed both high absolute detection values in the CLIP datasets and high conversion frequencies. Interestingly, some low-abundance miRNAs, such as hsa-miR-760 and hsa-miR-769-3p, also showed disproportionately high conversion levels, in line with our previous findings that uridines downstream of the seed sequence confer efficient crosslinking [14, 15].

We next applied the same analysis to our data derived from nuclear RBProximity-CLIP and nuclear Fractionation-CLIP (**Fig. 2G-H**). Using RBProximity-CLIP, no significant correlation was observed between miRNA expression and target site loading (ρ = 0.05, *p* = 0.48), in line with the lack of AGO2 in HEK293 cells nuclei. In contrast, the fractionation-derived nuclear dataset exhibited a clear positive correlation between detected target sites and miRNA expression levels (ρ = 0.39, *p* = 4.2 × 10⁻¹²), with many of the same miR-17∼92 members showing high expression and conversion values, likely reflecting the reduced specificity of biochemical fractionation. Fractionation-CLIP exhibited stronger correlations between miRNA abundance and seed-site conversion events in both cytoplasmic and nuclear fractions, likely due to the similar handling of total and fractionated extracts. RBProximity-CLIP, which includes additional processing steps, achieves higher specificity and retains biological signal (Spearman’s ρ = 0.19), albeit with reduced sensitivity.

AGO proteins are known to exhibit a distinct crosslinking pattern, with a preference for T-to-C conversion directly around the seed match [14, 15]. We therefore investigated T-to-C conversion events distribution along target sites of the ten top expressed miRNAs, including hsa-miR-19b-3p, hsa-miR-20a-5p and hsa-miR-16-5p (**Suppl. Fig. 1A**). As shown, the aggregated T-to-C conversion profile recapitulated strong enrichment immediately downstream of the miRNA seed sequences in cytoplasmic-but not in nuclear-RBProximity-CLIP cells (**Suppl. Fig. 1B, Suppl. Fig. 2**), suggesting capture of authentic cytoplasmic miRNA binding sites with RBProximity-CLIP, and minimal detection of background signal with nuclear RBProximity-CLIP. Taken together, we concluded that RBProximity-CLIP can capture functional AGO2–miRNA interactions.

### Targeting cytoplasmic, nuclear, and nucleolar YBX1–RNA interactions in HEK293 cells

We next aimed to demonstrate the capability of RBProximity-CLIP to investigate the subcellular RNA target profile of untagged, endogenous YBX1. YBX1 is a highly conserved, abundant, cold-shock domain-containing RBP, residing in both the nucleus and the cytoplasm [49]. The fact that YBX1 is implicated in several cellular processes, including RNA metabolism and transport [50], suggests its distinct locations may comprise separate functional subpopulations. Using cell imaging and biochemical fractionation in HEK293 cells, we could show YBX1’s nucleo-cytoplasmic distribution (**Fig. 3A,B**). Both approaches indicated a roughly 80%-20% distribution of YBX1 between the cytoplasm and the nucleus, respectively. Once we established that we could capture and process endogenous YBX1 RNPs (**Fig. 3C**), we proceeded to explore subcellular YBX1-RNA interactions using RBProximity-CLIP. Following a report of YBX1 involvement in nucleolar biology [51], we also targeted the YBX1 subpopulation in the nucleolus by generating a cell line that fused APEX2 to NIK3x, thereby targeting APEX2 to the nucleolus (**Fig. 3D,E**).

**Figure 3.**
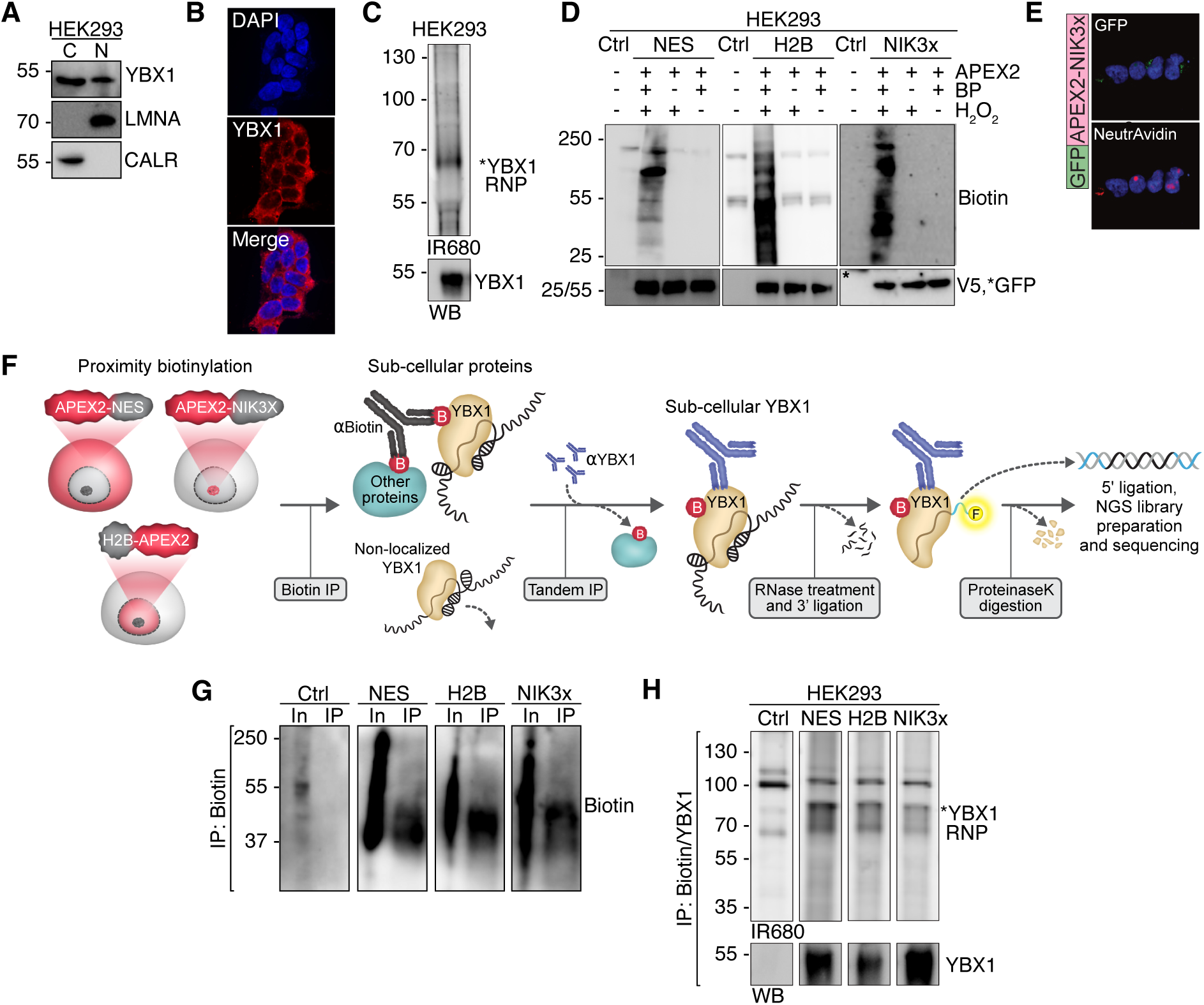
Capture of YBX1 subcompartmental RNA interactome. **(A)** YBX1, Lamin A/C (LMNA) and Calreticulin (CALR) Western blot analysis of biochemically fractioned HEK293 cells cytoplasmic (C) and nuclear (N) preparations. **(B)** YBX1 (red) immunofluorescence imaging in HEK293 cells co-stained with DAPI (blue). **(C)** Fluorescently imaged SDS-PAGE of 3′-ligated YBX1-RNA IP preparation (top) and a corresponding YBX1 Western blot analysis (bottom). **(D)** Streptavidin-HRP and V5 or GFP Western blot analysis of HEK293 whole cell lysates in the presence or absence of proximity biotinylation components: V5-APEX2–NES (NES), V5-APEX2–H2B (H2B) or GFP-APEX2–NIK3x (NIK3x), doxycycline induced expression, biotin-phenol (BP) and hydrogen peroxide (H_2_O_2_). V5-APEX2–NES migrates at approximately 25 kDa and V5-APEX2–H2B and GFP-APEX2–NIK3x migrates at approximately 55 or 54 kDa, respectively. **(E)** Fluorescent imaging of GFP-APEX2–NIK3x expressing HEK293 cells (green), co-stained with fluorescent NeutrAvidin (red) and DAPI (blue). **(F)** Schematic overview of the RBProximity-CLIP workflow. HEK293 cells expressing V5-APEX2–NES, V5-APEX2–H2B or GFP-APEX2–NIK3x undergo proximity biotinylation to label proteins in the cytoplasmic, nuclear or nucleolar compartments, respectively. Biotinylated proteins are immunoprecipitated using an anti-biotin antibody, followed by a sequential YBX1 IP in tandem. After RNase treatment and 3′ ligation of a fluorescent adapter, immobilized YBX1–RNA complexes are digested with proteinase K to release the crosslinked 3’-ligated RNA, which is then subjected to 5′ adapter ligation and downstream processing for next-generation sequencing (NGS) library preparation and sequencing. **(G)** Biotin western blot analysis of Input (In) and anti-biotin immunoprecipitated material from HEK293 cells stably expressing V5-APEX2–NES (NES), V5-APEX2–H2B (H2B), or GFP-APEX2–NIK3x (NIK3x), after proximity biotinylation. **(H)** Fluorescently imaged SDS-PAGE of 3′-ligated YBX1-RNA complexes (top) and a corresponding YBX1 western blot analysis (bottom). Complexes were sequentially captured using anti-biotin and anti-YBX1from HEK293 cell extracts after cytoplasmic (V5-APEX2–NES; NES), nuclear (V5-APEX2–H2B; H2B), or nucleolar (GFP-APEX2–NIK3x; NIK3x) proximity biotinylation.

To avoid RNP reconstitution after SDS-PAGE, we used a modified RBProximity-CLIP protocol for YBX1 (**Fig. 3F**). First, compartment-specific proteins were immobilized using immunoprecipitation with an anti-biotin antibody (**Fig. 3G**), followed by acetic acid elution, pH neutralization, and a sequential RBP-specific IP, isolating localized YBX1 RNPs. Following both IP procedures, immobilized YBX1-RNPs are exposed to RNase digestion and 3’ adaptor ligation, which enables visualization of the eluted RNP after SDS-PAGE (**Fig. 3H**). Then, guided by the 3’-adapter fluorescence to detect the size-shifted ligated RNP, compartment-specific YBX1-RNPs are excised, and the bound RNA fragments are released by Proteinase K digestion and further processed into a small RNA cDNA libraries for NGS analysis.

### YBX1 RBProximity-CLIP reveals conserved motif recognition and compartment-specific RNA binding patterns

We prepared three biological replicates of cytoplasmic, nuclear, and nucleolar RBProximity-CLIP datasets, along with a set of whole-cell YBX1 fPAR-CLIP libraries for comparative analysis. Pearson’s correlation indicated the strongest correlations between replicates (**Fig. 4A**); therefore, we merged the raw data of replicates for more powerful downstream analyses. All RBProximity-CLIP samples correlated well with the whole-cell fPAR-CLIP dataset and less well with each other (**Fig. 4B**), suggesting that compartment-distinct YBX1 interacted with unique RNA targets.

**Figure 4.**
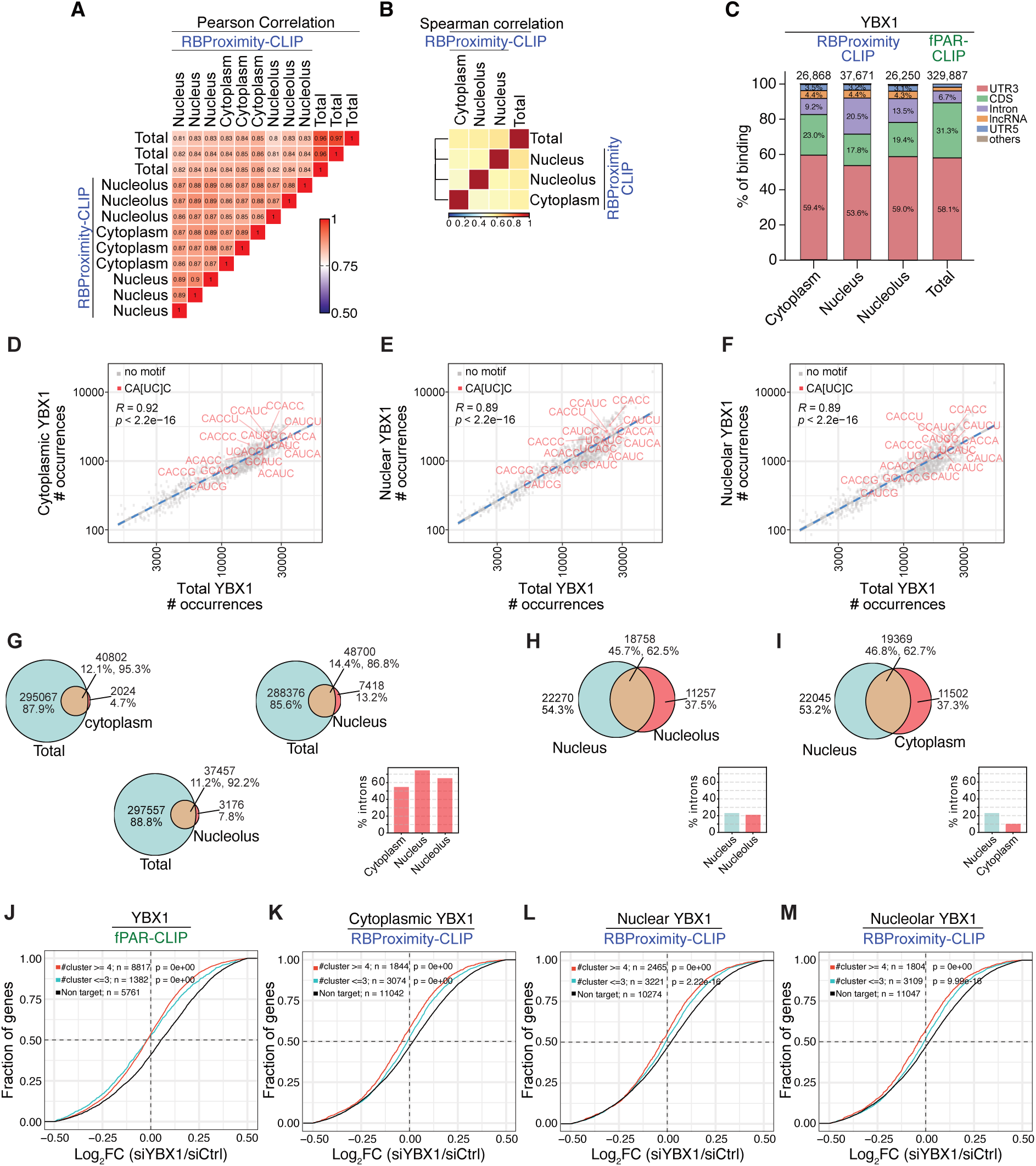
Comparative assessment of YBX1–RNA Interactions Across Subcellular Compartments. (A) Pearson correlation heatmap of transcript abundance across RBProximity-CLIP and whole-cell fPAR-CLIP datasets. **(B)** Spearman correlation heatmap of transcript abundance across RBProximity-CLIP and whole-cell fPAR-CLIP datasets, after pooling of replicates. Hierarchical clustering demonstrates the relationship between sample types. **(C)** Distribution of YBX1 binding between transcriptome annotations, obtained by cytoplasmic (V5-APEX2–NES), nuclear (V5-APEX2–H2B) and nucleolar (GFP-APEX2–NIK3x) RBProximity-CLIP and whole-cell fPAR-CLIP. Coding sequences (CDS), 3′ untranslated regions (UTR3), 5′ untranslated regions (UTR5), introns, and all other annotations. **(D)** Venn diagrams intersecting YBX1-bound RNA clusters identified by RBProximity-CLIP per compartment (cytoplasmic; V5-APEX2–NES, nuclear; V5-APEX2–H2B, and nucleolar; GFP-APEX2–NIK3x) and by whole-cell fPAR-CLIP, number and percentage of overlapping and unique clusters are displayed. For the clusters identified exclusively by RBPoximity-CLIP (red), percentage of intronic clusters is displayed in the bottom-right bar graph. **(E,F)** Venn diagrams intersecting YBX1-bound RNA clusters in the nucleus (V5-APEX2–H2B) versus in the nucleolus (GFP-APEX2–NIK3x) (**E**) or the cytoplasm (**F**), as identified by RBProximity-CLIP. Number and percentage of overlapping and unique clusters are displayed. For compartment-exclusive clusters (red), percentage of intronic clusters is displayed in the bottom-right bar graph. **(G-I)** 5-mer frequency among YBX1 clusters detected by whole-cell fPAR-CLIP, relative to its frequency among YBX1 cytoplasmic (V5-APEX2–NES) (**G**), nuclear (V5-APEX2–H2B) (**H**) and nucleolar (GFP-APEX2–NIK3x) (**I**) clusters obtained by RBProximity-CLIP. Motifs containing the CA[UC]C consensus sequence are labelled (orange). A linear regression line (blue) is shown with correlation coefficient. **(J-M)** Cumulative distribution of total-cell (**J**), cytoplasmic (V5-APEX2–NES) (**K**), nuclear (V5-APEX2–H2B) (**L**) and nucleolar (GFP-APEX2–NIK3x) (**M**) YBX1 gene targets along stratified differential expression observed upon YBX1 knockdown (log₂ fold-change [siYBX1/siCtrl]). Targets are segmented by number of clusters per gene (red≥4 YBX1, cyan ≤3), and the distribution of expressed non-YBX1 targets is also presented (black). Statistical significance was assessed using Kolmogorov-Smirnov (K-S) test.

Basic filtration of data included the following criteria: (i) False Discovery Rate (FDR) ≤ 0.05, (ii) read depth per cluster (binding site) ≥ 10, (iii) cluster length ≥ 10 nt, (iv) T-to-C conversion locations per cluster ≥ 2. Across all conditions, the total number of clusters decreased progressively with each applied filter, reflecting increased stringency, with 26,000-38,000 RBProximity-CLIP clusters per subcellular compartment and 330,000 clusters per whole-cell fPAR-CLIP passing our filters (**Fig. 4C, Suppl. Fig. 3A**). As observed for AGO2, a reduced number of detected clusters following RBProximity-CLIP relative to canonical whole-cell fPAR-CLIP probably represents RBProximity-CLIP specificity, along with reduced sensitivity due to additional processing.

Given YBX1’s distribution to both the cytoplasm, which contains mature mRNAs, and the nucleus, which contains pre-mRNAs, we first compared YBX1 occupancy along functional transcript elements between the different compartments (**Fig. 4C**). Generally, YBX1 binding along mature mRNA elements (3’UTR, CDS, and 5’UTR), and pre-mRNAs (introns), was similar across samples, with dominant 3’UTR binding (53-59% of clusters), consistent with previously published studies [52, 53]. With 80% of YBX1 observed in the cytoplasm (**Fig. 4A,B**), we expected total cell YBX1 binding trends to be most similar to those obtained by cytoplasmic RBProximity-CLIP. Indeed, these two samples comprised the lowest portion of intronic clusters, while nuclear RBProximity-CLIP showed a 50% higher intronic occupancy, suggesting that RBProximity-CLIP indeed enriches for YBX1 binding sites from the targeted compartments (**Fig. 4C**).

To investigate YBX1 binding preference across compartments, we computed the relative abundance of 5-mers between the RBProximity-CLIP and fPAR-CLIP libraries (**Fig. 4D-F**), and relative to generated controls (**Suppl. Fig. 3B-E**). As shown, all RBProximity-CLIP datasets and whole-cell fPAR-CLIP consistently present enrichment of the previously described YBX1 binding consensus motif – CA[UC]C [54]. This analysis confirms that RBProximity-CLIP drives detection of authentic YBX1-RNA interactions, indicating that YBX1 molecular binding preference remains spatially consistent, while target specificity is most probably set by the localized context and the availability of targets.

Next, we used the binding coordinates of clusters to define cross-compartment shared and exclusive YBX1 targets. Almost all compartment-exclusive YBX1-targets were also detected by whole-cell fPAR-CLIP analysis (**Fig. 4G**), indicating that it captured nearly the full spectrum of YBX1-bound RNAs across all subcellular compartments. The small portions of compartment-exclusive targets that were not detected in whole-cell fPAR-CLIP were primarily intronic, likely reflecting the scarcity of intronic transcripts within cells. Notably, also among these target sets, the intronic content was relatively reduced in samples obtained by cytoplasmic RBProximity-CLIP (**Fig. 4G, bar plot).** A sub-nuclear comparison shows that ∼37% of targets detected by nucleolar RBProximity-CLIP were exclusive, not detected by whole-nucleus RBProximity-CLIP (**Fig. 4H**). Comparing nuclear and cytoplasmic RBProximity-CLIP datasets revealed a 4-fold higher frequency of intron binding among nucleus-specific YBX1 targets (**Fig. 4I**). The specificity obtained by RBProximity-CLIP is also demonstrated by comparing RBProximity-CLIP signal along specific binding sites. For example, an intronic binding site within the SFPQ pre-mRNA is detected in total cells as well as nuclear and nucleolar RBProximity-CLIP, but undetected by cytoplasmic RBProximity-CLIP (**Suppl. Fig. 3F**). Similarly, the snoRNA, SNORD3B is also specifically bound by non-cytoplasmic YBX1, possibly in accordance with a report attributing nucleolar localization for YBX1 [51] (**Suppl. Fig. 3G**).

Finally, we tested whether RBProximity-CLIP enriched for functional YBX1 binding sites and profiled the transcriptome after knockdown of YBX1 with specific siRNAs (**Suppl. Fig. 3H,I**). Our results show that YBX1 targets tended to decrease in abundance after YBX1 knockdown (**Fig. 4J-M**), suggesting that independent of the profiled compartment, RBProximity-CLIP captured functionally relevant YBX1-RNA interactions.

### Targeting cytoplasmic, nuclear, and nucleolar ELAVL1–RNA interactions in HEK293 cells

To further evaluate the versatility of RBProximity-CLIP, we targeted another endogenous abundant RBP, HuR/ELAVL1. Unlike YBX1, ELAVL1 primarily localizes to the nucleus but can shuttle to the cytoplasm [55], as confirmed by nucleocytoplasmic fractionation followed by Western blot analysis and immunofluorescence imaging (**Fig. 5A,B**). We utilized our established HEK293 cell lines expressing localized APEX2 to dissect the RNA interactome of endogenous ELAVL1 across the cytoplasm, nucleus, and nucleolus (**Fig. 5C,D**). After confirming the correlation between biological replicates (**Fig. 5E**), datasets were merged to increase power for downstream analyses. Hierarchical clustering of the merged datasets revealed that RBProximity-CLIP of nuclear ELAVL1 showed the highest similarity to total cell ELAVL1 fPAR-CLIP, consistent with the predominantly nuclear localization of ELAVL1 (**Fig. 5F**).

**Figure 5.**
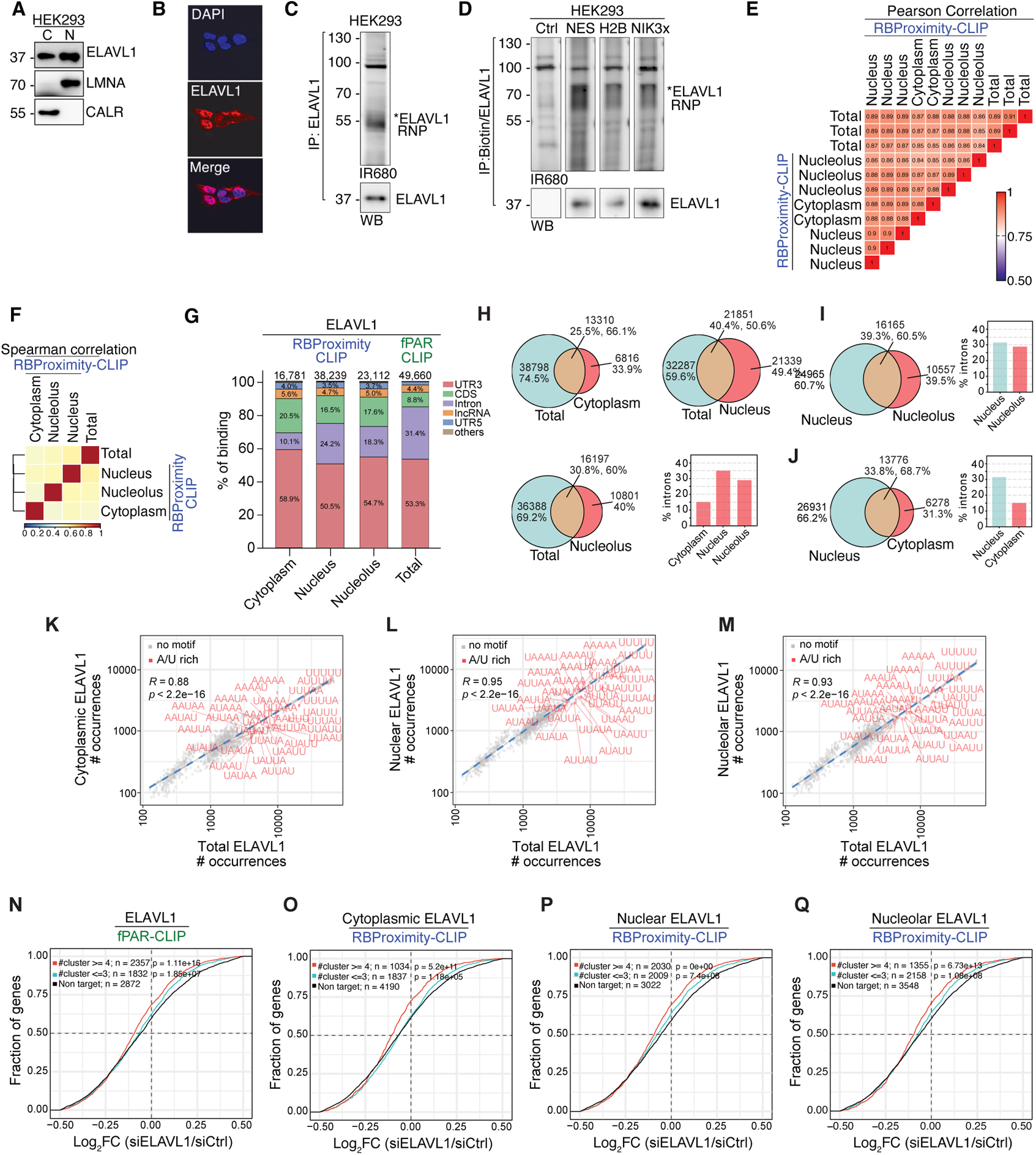
Subcompartmental RNA interactome of ELAVL1. **(A)** ELAVL1, Lamin A/C (LMNA) and Calreticulin (CALR) Western blot analysis of biochemically fractioned HEK293 cells cytoplasmic (C) and nuclear (N) preparations. **(B)** ELAVL1 (red) immunofluorescence imaging in HEK293 cells co-stained with DAPI (blue). **(C)** Fluorescently imaged SDS-PAGE of 3′-ligated ELAVL1-RNA whole cell IP preparation (top) and a corresponding ELAVL1 Western blot analysis (bottom). **(D)** Fluorescently imaged SDS-PAGE of 3′-ligated ELAVL1-RNA complexes (top) and a corresponding Western blot analysis (bottom). Loaded complexes were sequentially captured using anti-biotin and anti-ELAVL1 from HEK293 cell extracts after cytoplasmic (V5-APEX2–NES; NES), nuclear (V5-APEX2–H2B; H2B), or nucleolar (GFP-APEX2–NIK3x; NIK3x) proximity biotinylation. **(E)** Pearson correlation heatmap of transcript abundance across RBProximity-CLIP and whole-cell fPAR-CLIP datasets. **(F)** Spearman correlation heatmap of transcript abundance across RBProximity-CLIP and whole-cell fPAR-CLIP datasets, after pooling of replicates. Hierarchical clustering demonstrates the relationship between sample types. **(G)** Distribution of ELAVL1 binding between transcriptome annotations, obtained by cytoplasmic (V5-APEX2–NES), nuclear (V5-APEX2–H2B) and nucleolar (GFP-APEX2–NIK3x) RBProximity-CLIP and whole-cell fPAR-CLIP. Coding sequences (CDS), 3′ untranslated regions (UTR3), 5′ untranslated regions (UTR5), introns, and all other annotations. **(H)** Venn diagrams intersecting ELAVL1-bound RNA clusters identified by RBProximity-CLIP per compartment (cytoplasmic; V5-APEX2–NES, nuclear; V5-APEX2–H2B, and nucleolar; GFP-APEX2–NIK3x) and by whole-cell fPAR-CLIP, number and percentage of overlapping and unique clusters are displayed. For the clusters identified exclusively by RBPoximity-CLIP (red), percentage of intronic clusters is displayed (bottom-right bar graph). **(I-J)** Venn diagrams intersecting ELAVL1-bound RNA clusters in the nucleus (V5-APEX2–H2B) versus in the nucleolus (GFP-APEX2–NIK3x) (**I**) or the cytoplasm (V5-APEX2–NES) (**J**), as identified by RBProximity-CLIP. Number and percentage of overlapping and unique clusters are displayed. For compartment-exclusive clusters (red), percentage of intronic clusters is displayed in the bottom-right bar graph. **(K-M)** 5-mer frequency among ELAVL1 clusters detected by whole-cell fPAR-CLIP, relative to its frequency among YBX1 cytoplasmic (V5-APEX2–NES) (**K**), nuclear (V5-APEX2–H2B) (**L**) and nucleolar (GFP-APEX2–NIK3x) (**M**) clusters obtained by RBProximity-CLIP. Motifs containing the CA[UC]C consensus sequence are labelled (orange). A linear regression line (blue) is shown with correlation coefficient. **(N-Q)** Cumulative distribution of whole-cell (**N**), cytoplasmic (V5-APEX2–NES) (**O**), nuclear (V5-APEX2–H2B) (**P**) and nucleolar (GFP-APEX2–NIK3x) (**Q**) ELAVL1 gene targets along stratified differential expression observed upon ELAVL1 knockdown (log₂ fold-change [siELAVL1/siCtrl]). Targets are segmented by number of clusters per gene (red≥4 ELAVL1, cyan ≤3), and the distribution of expressed non-ELAVL1 targets is also presented (black). Statistical significance was assessed using Kolmogorov-Smirnov (K-S) test.

We stringently filtered our obtained data as described above for YBX1 and again observed reduced RBProximity-CLIP sensitivity relative to whole-cell fPAR-CLIP, with 16,000-38,000 clusters detected per subcellular compartment compared to 49,000 clusters passing filtration in whole-cell fPAR-CLIP (**Fig. 5G, Suppl. Fig 4A**). The distribution of ELAVL1-bound RNA clusters across genomic regions aligned with ELAVL1’s predominant nuclear localization (**Fig. 5G**), and accordingly, nuclear RBProximity-CLIP yielded the highest number of ELAVL1-bound clusters (38,239), enriched for intronic elements, compared to cytoplasmic RBProximity-CLIP (**Fig. 5G**).

Using binding site coordinates to identify compartment-specific ELAVL1 clusters identified subsets of unique clusters (**Fig. 5H-J**), reflecting the compartment of origin. Analysis of ELAVL1-bound sequence motifs across compartments (**Fig. 5K-M, Suppl. Fig 4B-E**) confirmed that ELAVL1 bound A/U-rich elements as previously reported [56] across all assayed compartments, suggesting that target availability is likely the main driver of compartment-specific differences in the interactome. The specificity obtained by RBProximity-CLIP is also demonstrated by comparing RBProximity-CLIP signal along specific binding sites. For example, intronic binding sites along NUCKS1 or SRRM2 genes are exclusively undetected cytoplasmic RBProximity-CLIP (**Suppl. Fig 4F,G**).

Finally, to assess recovery of functional ELAVL1 targets by RBProximity-CLIP we integrated our data with available ELAVL1 loss-of-function transcriptome profiles [56]. ELAVL1 targets, from any subcellular compartment, showed decreased abundance upon ELAVL1 depletion (**Fig. 5N-Q**), in line with capture of functional binding sites. However, in light of the large overlap between ELAVL1 targets across compartments, this does not necessarily imply that ELAVL1 regulated mRNA abundance in all compartments. Further investigation is necessary to understand the precise molecular function of ELAVL1 in the nucleolus or cytoplasm, which likely differs from its reported role in splicing regulation.

## Discussion

In this study, we developed RBProximity-CLIP, combining proximity-biotinylation and UV-induced RBP-RNA crosslinking, followed by sequential RBP- and biotin-affinity purifications, to enable subcellularly resolved mapping of RBP-RNA interactions. RBProximity integrates fPAR-CLIP [14, 42] with Proximity-CLIP [10, 40], while developing procedures for sequential affinity purification of RBP-RNA complexes, including reconstitution of a gel-extracted RNP, the sequence of RBP IP and anti-biotin IP with streptavidin affinity purification, and adaptation of on- and off-beads molecular manipulations that are required for small RNA cDNA NGS library preparation. A detailed and elaborate characterization of intermediate biochemical and molecular products, benchmark comparison to biochemical fractionation, and data analyses of the obtained results, confirm RBProximity-CLIP specificity, while also shedding light on the sensitivity cost involving RBProximity-CLIP. Collectively, by providing high specificity and adequately sensitive detection, RBProximity-CLIP enabled spatial profiling of RNA targets for three distinct RBPs: AGO2, YBX1, and ELAVL1, within the cytoplasmic, nuclear, and nucleolar compartments.

CLIP methods were, in several cases, coupled with biochemical fractionation to characterize nucleocytoplasmic and chromatin-associated RBP targets [16, 41, 57, 58]; however, these approaches are often limited to membrane-enclosed compartments, are labor-intensive, and usually suffer from cross-contamination from other cellular compartments due to incomplete separation. By contrast, APEX2-based RBProximity-CLIP offers several advantages: 1) not limited to fractionable compartments, 2) less cell extract manipulations compared to biochemical fractionation, 3) reduced background, 4) rapid labelling and thus temporal specificity for capture of dynamic processes such as triggered stress granule formation [41], infection, differentiation, cell cycle, etc.

Recently, Colocalization CLIP (coCLIP) [41] was introduced and is an analogues method to RBProximity-CLIP. Using coCLIP, subcellular ELAVL1-RNA interactions were characterized during cell stress. While conceptually similar, important differences provide RBProximity-CLIP with increased specificity and sensitivity. First, RBProximity-CLIP relies on 4SU-dependent UVA/UVB induced protein-RNA crosslinking, which enables efficient crosslinking, as well as results in T-to-C conversion events indicative of authentic crosslinking events prior to cell lysis [14]. Furthermore, stringent size-selection using fluorescently labeled RNP purifications enables further reduction of background signals in RBProximity-CLIP. RBProximity-CLIP enables the use of either streptavidin affinity purification or anti-biotin immunoprecipitation, providing versatility that, in our experience, is beneficial for the flexible application of this method to various endogenous RBPs.

The major limitation of RBProximity-CLIP is the reduced sensitivity linked to the additional required processing. Nevertheless, our results indicate adequate detection by RBProximity-CLIP enabling discovery in the field of subcellular RBP functions. Another factor characterizing RBProximity-CLIP is the requirement for localized APEX2 expression, limiting transition between different cell lines, and essentially rendering RBProximity-CLIP irrelevant for fresh, non-genetically manipulated tissue.

Using AGO2, a well-characterized RBP central to RNA silencing, we validated RBProximity-CLIP’s capacity to recover canonical miRNA binding profiles from the nuclear or cytoplasmic compartments. The near-absence of nuclear clusters in HEK293 cells underscores the specificity of the approach and its advantage over fractionation-based CLIP, which often exhibits nuclear signal leakage [7].

RBProximity-CLIP targeting YBX1 and ELAVL1 provided complementary insights into the putative multifunctionality of these proteins across subcellular compartments. YBX1 exhibited dual cytoplasmic and nuclear RNA interactomes consistent with its multifunctional role in RNA metabolism, retaining its molecular recognition of the CA[UC]C motif across all compartments, indicating a conserved RNA recognition mode. Likewise, RBProximity-CLIP captured ELAVL1’s expected AU-rich binding motif. For both YBX1 and ELAVL1, RBProximity-CLIP captured both shared and compartment-specific interaction profiles, indicative of its discovery potential for compartment-specific RBP-RNA interactions.

An important factor affecting the interpretation of binding to pre-mRNA is the limited reproducibility for binding sites detection along introns, observed across CLIP approaches [9, 59]. The vast intronic sequence space, offering a multitude of binding sites for RBPs that recognize short, degenerate sequence elements, combined with the very short half-life of introns, limits the likelihood of binding events being resampled in replicates.

In summary, by studying the interaction of three RBPs with RNAs in three subcellular compartments, we demonstrate the utility of RBProximity-CLIP as an experimental framework for investigating RBP–RNA interactions at subcellular resolution and with temporal precision.

With the large availability of localized APEX2 constructs, RBProximity-CLIP is readily applicable for exploring RNA regulation at numerous subcellular locations.

## Materials and Methods

### APEX2 compatible cell line generation

HEK293-APEX2–NES and HEK293-APEX2–H2B cells were previously generated [10]. FLAG-HA-AGO2 expression was introduced into HEK293, HEK293-APEX2–NES and HEK293-APEX2–H2B cells by lentiviral transduction according to the protocol described by Tandon and colleagues [60]. Cells were co-transfected with plasmids encoding a mixture of viral packaging proteins VSV-G (Addgene, 12259), viral backbone psPAX2 plasmid (Addgene, 12260) and FLAG-HA-AGO2 (Addgene, 91978) at ratio 3:2:4, where the final plasmid concentration in the media was 2.25 μl/ml. Cells were kept in selection media [complete media supplemented with 2 μg/ml Puromycin (SigmaAldrich, P9620)] for 10 days, when control, none transduced, cells died. Following, cell clones were cultured in complete media.

The GFP-APEX2–NIK3x (Addgene, 129274) plasmid was used to clone the GFP-APEX2–NIK3x sequences into the pcDNA5/FRT/TO vector, producing pcDNA5-FRT-GFP-APEX2–NIK3x expression vectors. The inFusion system was used for cloning (Takara, 638909), all primers were designed using the Takara primer design system, and the constructs were verified by Sanger sequencing. Subsequently, the plasmid was introduced into HEK293-Flp-In-T-Rex cells (Thermofisher Scientific, R78007) using the Flp-In-T-Rex core kit standard manual (ThermoFisher, K650001), resulting in the generation of HEK293-APEX2–NIK3x stable cell line. The expression and subcellular localization of the APEX2 fusion constructs were validated by western blot and immunofluorescence assays, described below.

### Cell culture conditions

HEK293 (CRL-1573) were obtained from ATCC and cultured in DMEM media (Gibco, 11995065) supplemented with 10% Fetal Bovine Serum (FBS) (Gibco, 10270106) and 1% Penicillin-Streptomycin (P/S) solution (Gibco, 15140122). The media for HEK293-Flp-In-T-Rex cells was additionally supplemented with 200 µg/ml Zeocin (Gibco, R25005) and 5 µg/ml Blasticidin (Gibco, A1113903). HEK293-APEX2–NES, HEK293-APEX2–H2B, HEK293-APEX2–NIK3x cell lines were cultured in complete media supplemented with 100 μg/ml of Hygromycin B (Gibco, 10687010) and 5 µg/ml Blasticidin. All cells were cultured at 37°C, with 5% CO_2_.

### Proximity-dependent biotinylation

Cells were seeded at a density of 35,000 cells/cm^2^ 48 hours prior to the biotinylation reaction in complete media, without selection antibiotics. 16 hours before the reaction, the expression of APEX2 fusion constructs was induced with 2 μg/ml doxycycline (Fisher Scientific, BP2653-1). The next day, 500 mM biotin tyramide (Iris Biotech, 41994-02-09) was diluted to a concentration of 10 mM in warm complete media and added to the cells to a final concentration of 500 μM. Cells were incubated at 37°C for 30 minutes. Next, to trigger the biotinylation reaction, 100 mM H_2_O_2_ was added to the media to a final concentration of 1 mM. After precisely 60 seconds, the media was quickly poured off and the cells were washed three times with freshly prepared quenching solution [10 mM sodium ascorbate in water (Sigma Aldrich, A4034), 10 mM sodium azide in water (Sigma Aldrich, S2002) and 5 mM Trolox in DMSO (Sigma Aldrich, 238813) prepared in 1xPBS (Gibco, 10010023)]. Subsequently, cells were UV-crosslinked (See below), scraped into 1.5 ml tubes and either snap-frozen in liquid nitrogen or immediately subjected to cell lysis and protein extraction.

### Immunofluorescence imaging

HEK293 and HEK293-APEX2–NIK3x cell lines were plated at a density of 35,000 cells/cm^2^ on glass coverslips inside 6-well plates. For visualization of APEX2 protein fusions and biotinylated proteins, APEX2-compatible HEK293 cell lines were processed according to the biotinylation procedure described above and fixed with 4% paraformaldehyde (Solveco, 1267) for 15 minutes at room temperature. For visualization of YBX1 and ELAVL1 protein, HEK293 cells were fixed directly in 4% paraformaldehyde for 15 minutes at room temperature. Next, cells were washed three times with PBS and permeabilized in 0.1% Triton X-100 (Merck, 2341264) in deionized water for 5 minutes at room temperature. Subsequently, cells were washed three times with 1x PBS, blocked with 5% BSA (Fisher Scientific, BP9703-100) in 1x PBS for 1 hour at room temperature, and stained with primary antibodies overnight, as outlined in the table below. Following this, the cells were washed three times and incubated for 1 hour with secondary antibodies, as listed in the table below. GFP-APEX2–NIK3x fusion protein was visualized through GFP fluorescence. Following this, cells were washed twice with 1x PBS and incubated with 300 nM 4′,6-diamidino-2-phenylindole (DAPI; ThermoFisher, D1306) for 10 minutes at room temperature to visualize the nuclei. Slides were mounted with ProLong Diamond Antifade Mountant (ThermoFisher, P36961). Confocal images were acquired with Zeiss LSM780 and processed with ImageJ software.

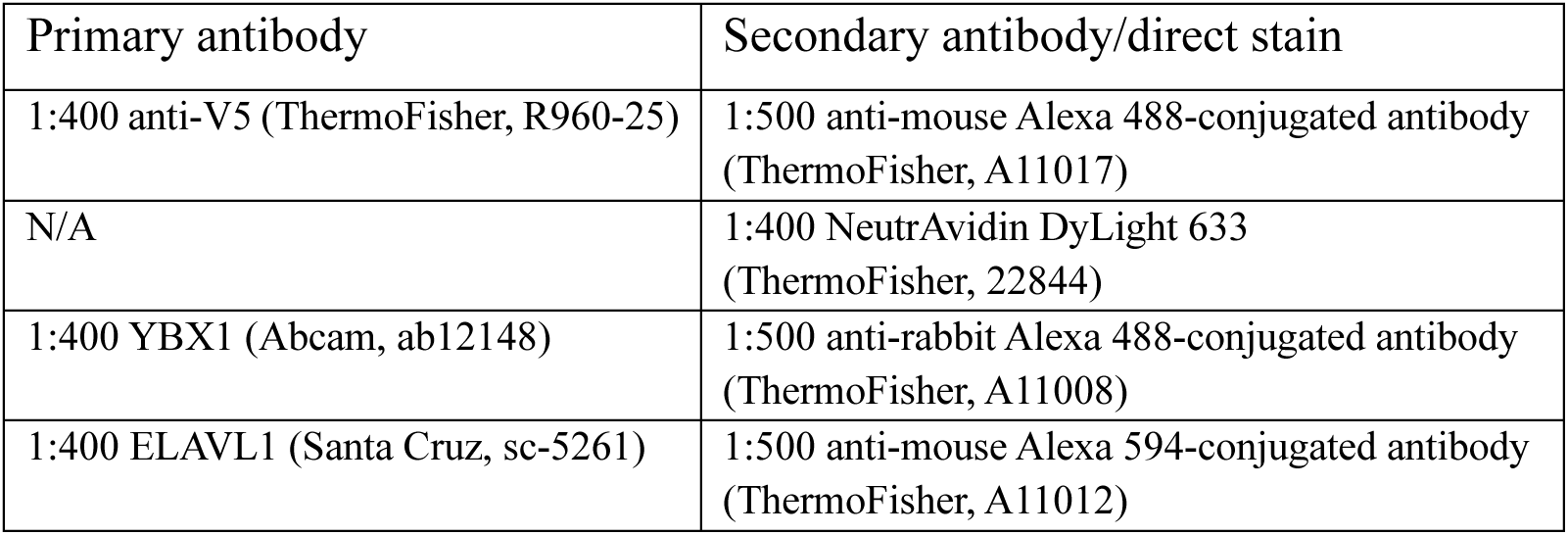

| Primary antibody | Secondary antibody/direct stain |
| --- | --- |
| 1:400 anti-V5 (ThermoFisher, R960-25) | 1:500 anti-mouse Alexa 488-conjugated antibody (ThermoFisher, A11017) |
| N/A | 1:400 NeutrAvidin DyLight 633 (ThermoFisher, 22844) |
| 1:400 YBX1 (Abcam, ab12148) | 1:500 anti-rabbit Alexa 488-conjugated antibody (ThermoFisher, A11008) |
| 1:400 ELAVL1 (Santa Cruz, sc-5261) | 1:500 anti-mouse Alexa 594-conjugated antibody (ThermoFisher, A11012) |

### Cell lysis

Cells were lysed in RIPA buffer [50 mM Tris, 150 mM NaCl, 0.1% (wt/vol) SDS, 0.5% (wt/vol) sodium deoxycholate, and 1% (vol/vol) Triton X-100, pH 7.5], supplemented with protease and phosphatase inhibitor cocktail (ThermoFisher, 78445), incubated on ice for 15 minutes, and extracts were cleared by centrifugation at 21,000 *g* for 15 minutes at 4 °C. Protein concentration was determined with Pierce 660-nm assay (ThermoFisher, 22660). For cells lysed after proximity-biotinylation RIPA buffer was additionally supplemented with fresh quenchers (10 mM sodium azide, 10 mM sodium ascorbate, and 5 mM Trolox).

### Biochemical fractionation

Cells were fractionated as described in [21, 61]. Briefly, cell pellets were dissolved gently in Hypotonic Lysis Buffer (HLB) [10 mM Tris–HCl, pH 7.5; 10 mM NaCl; 3 mM MgCl2; 0.3% NP-40 (v/v); and 10% glycerol (v/v)], supplemented with protease and phosphatase inhibitor cocktail and incubated for 5 minutes on ice. Next, the nuclei were isolated by centrifugation at 200 *g* for 2 minutes at 4 °C and the supernatants were collected as the cytoplasmic fraction. The pellet, nuclear fraction, was further washed three times in HLB, and each time, it was isolated by centrifugation at 200 *g* for 2 minutes at 4 °C. The washed nuclear pellet was dissolved with Nuclear Lysis Buffer (NLB) [20 mM Tris–HCl, pH 7.5; 150 mM KCl; 3 mM MgCl_2_; 0.3% NP-40 (v/v), and 10% glycerol (v/v)], supplemented with protease and phosphatase inhibitor cocktail at the same volume as the corresponding cytoplasmic fraction. The nuclear fraction was sonicated twice, with 10 seconds on and 30 seconds off, at 60% amplitude (Sonics, VCX130). Both cytoplasmic and nuclear fractions were cleared by centrifugation at 21,000 *g* for 15 minutes at 4 °C.

### SDS page and immunoblotting

Western blot experiments were performed as described earlier [61]. Briefly, 10–20 μg of protein was used for both whole cell lysates and the cytoplasmic fractions. The nuclear fractions were loaded at 1:1 volume ratio with the corresponding cytoplasmic fractions. Samples were loaded to 8%, 10% or 12% acrylamide:bisacrylamide SDS PAGE gels and, after electrophoresis, the proteins were transferred onto Nitrocellulose membrane (Cytiva, 1060000) and the blots stained with Ponceau S, washed, and blocked with 3 % non-fat dry milk (Sigma Aldrich, 70166) in 0.1 % Tween-20 in TBS (TBS-T). Membranes were incubated with primary antibodies, or for the detection of biotinylated proteins, HRP-conjugated streptavidin, at the dilutions and times indicated in the table below. Following the incubation, the membrane was washed and incubated with 1:3000 HRP-conjugated anti-rabbit IgG (Cytiva, NA934) or anti-mouse IgG (Cytiva, NA931) in 1% non-fat dry milk for 1 hour at room temperature and further washed in TBS-T. The signal was detected by chemiluminescence using ThermoScientific SuperSignal™ West Dura Extended Duration Substrate (ThermoFisher, 34076).

| Antibody | Catalog number | Dilution | Incubation |
| --- | --- | --- | --- |
| AGO2 | Abcam, ab32381 | 1:4000 in 1% milk in TBS-T | Overnight |
| Lamin A/C | Santa Cruz<br>sc-376248HRP | 1:5000 in 1% milk in TBS-T | 1 hour |
| Streptavidin-HRP | Molecular Probes, S911 | 1:2000 in 1% Bovine Serum Albumin (BSA, BP9703, Fisher Bioreagents) in TBS-T | 2 hours |
| V5 | ThermoFisher, R960-25 | 1:2000 in 1% BSA in TBS-T | 1 hour |
| Calreticulin | Santa Cruz<br>sc-373863HRP | 1:1000 in 1% milk in TBS-T | 1 hour |
| FLAG | Sigma Aldrich, F1804 | 1:5000 in 1% milk in TBS-T | Overnight |
| YBX1 | Abcam, ab12148 | 1:5000 in 1% milk in TBS-T | Overnight |
| ELAVL1 | Santa Cruz, sc-5261 | 1: 1000 in 1% milk in TBS-T | Overnight |

### YBX1 silencing and RNA sequencing

About 6×10^5^ cells were seeded in 60-mm plates and transfected the next day with 50 pmol of siRNA targeting YBX1 (ThermoFisher Scientific, 4390824, siRNA ID s9731) or a control scrambled siRNA (ThermoFisher Scientific, 4390843) preincubated for 20 min at room temperature with 5 µL of Lipofectamine™ RNAiMAX transfection reagent (ThermoFisher Scientific, 13778150) in OPTI-MEM medium. Cells were collected 72 hours after transfection. Total protein was extracted from half of the cells using RIPA buffer and analyzed by Western blot to assess silencing efficiency. Total RNA was isolated from the remaining cells using TRIzol™ reagent (ThermoFisher Scientific, 15596018) and treated with DNAse I using the Direct-zol RNA Miniprep kit (Zymo research, R2052) according to the manufacturer’s instructions. Each siRNA transfection was performed in biological triplicate.

rRNA-depleted libraries for RNA-sequencing were generated from 1 µg of total RNA using the NEBNext® rRNA Depletion Kit v2 (New England Biolabs, E7400X) and NEBNext® Ultra™ II Directional RNA Library Prep Kit for Illumina® (New England Biolabs, E7760L) according to the manufacturer’s instructions. The quality of the initial RNA material and of the final cDNA libraries was assessed using the TapeStation system (Agilent Technologies). Final libraries were sequenced on an Illumina NovaSeq X Plus platform (Illumina) to generate paired-end reads (2 × 50 bp).

Sequencing quality was assessed using *FastQC* (v0.11.8). Reads were aligned to the human reference genome (hg38) with *STAR* (v2.7.9a) using default parameters and the option --quantMode GeneCounts to obtain raw read counts per gene, based on the ncbiRefSeq annotation. Differential expression analyses were conducted with *R* (v4.4.2) and *DESeq2* (v1.46.0) using median-of-ratios normalized read counts and Benjamini-Hochberg adjusted p-values. Genes were considered differentially expressed based on the following criteria: baseMean > 10, adjusted p-value < 0.05 and log_2_(fold change[treated/control]) > 1 (up-regulated) or < -1 (down-regulated).

### 4-thiouridine labelling and UV crosslinking

For the fPAR-CLIP and RBProximity-CLIP procedures described below, cells were metabolically labelled with 4-thiouridine (4SU, Carbosynth, NT06186) and UV-crosslinked. 4SU was added directly to the growth media to a final concentration of 100 µM for 16 hours. APEX2-expressing and control HEK293 cell lines were then subjected to proximity biotinylation as described above. Next, cells were washed once with 1xPBS and UV-crosslinked at 312-nm at a dose of 0.15 J/cm^2^ for 5 minutes. Cells were collected by scraping.

### Whole-cell Fluorescent PhotoActivatable Ribonucleoside-enhanced CrossLinking and ImmunoPrecipitation (fPAR-CLIP)

HEK293 cells were seeded at a density of 35,000 cells/cm^2^ and collected after 48 hours following 4SU labelling and UV crosslinking. YBX1 and ELAVL1 fPAR-CLIP were performed targeting endogenous proteins using 4 µg antibodies against YBX1 and ELAVL1, respectively, coupled to 20 µl of Protein G beads magnetic beads from 1 mg of whole cell lysate. Following fPAR-CLIP library preparation, sequencing, and initial data processing were performed as described in [42]. Briefly, unprotected RNA was digested on beads with 50 μl of 1 U/μl RNase T1 (ThermoFisher, EN0541) for 15 minutes at room temperature with shaking at 800 rpm. Following, the beads were washed three times with RIPA buffer and once with dephosphorylation buffer (50 mM Tris–HCl, pH 7.5, 100 mM NaCl, 10 mM MgCl_2_). Next, the RNA was dephosphorylated with 30 μl of 0.5 U/μl Quick CIP (New England Biolabs, M0525S) for 10 minutes at 37°C with shaking at 800 rpm. Next, the beads were washed three times with dephosphorylation buffer and three times with ligation buffer (50 mM Tris-HCl, pH 7.5, 10 mM MgCl_2_). Following, 0.5 μM fluorescent 3′ adapter (MultiplexDX, MDX-O-226; see **Table 1** below) was ligated with T4 Rnl2(1–249)K227Q (New England Biolabs, M0351) overnight at 4°C with shaking and washed three times with PNK/ligation buffer. After washing, RNA was phosphorylated using T4 PNK (New England Biolabs, M0201S) for 30 minutes at 37°C with shaking at 800 rpm and washed three times with RIPA buffer. To elute the proteins, the beads were incubated at 95°C for 10 minutes in 40 μl of 3 × SDS Laemmli buffer. Next, the eluates were separated on a 4–12% SDS/PAGE gels and visualized on the IR680 channel (Chemidoc MP system, Bio Rad). Subsequently, YBX1-RNP bands (around 80 kDa) and ELAVL1-RNP bands (around 70 kDa) were cut out from the gel with a sterile blade, proteins digested with Proteinase K (Sigma Aldrich, RPROTK-RO), and released RNA isolated via phenol:chloroform phase separation. Following, 5′ adapter ligation was performed on the purified RNA samples with 0.5 μM of the adapter (MultiplexDX, MDX-O-264) and Rnl1 T4 RNA ligase (ThermoFisher, EL0021) for 1 hour at 37°C. Next, the RNA was reverse transcribed using SuperScript IV Reverse Transcriptase (ThermoFisher, 18090010) according to the manufacturer’s instructions. The libraries were amplified in a series of PCR reactions performed using Platinum Taq DNA polymerase (ThermoFisher, 10966034) and size selected with 3% Pippin Prep (Sage Science, CSD3010). Sequencing of the libraries was carried out on the Illumina NovaSeq 6000 platform.

**Table 1.**
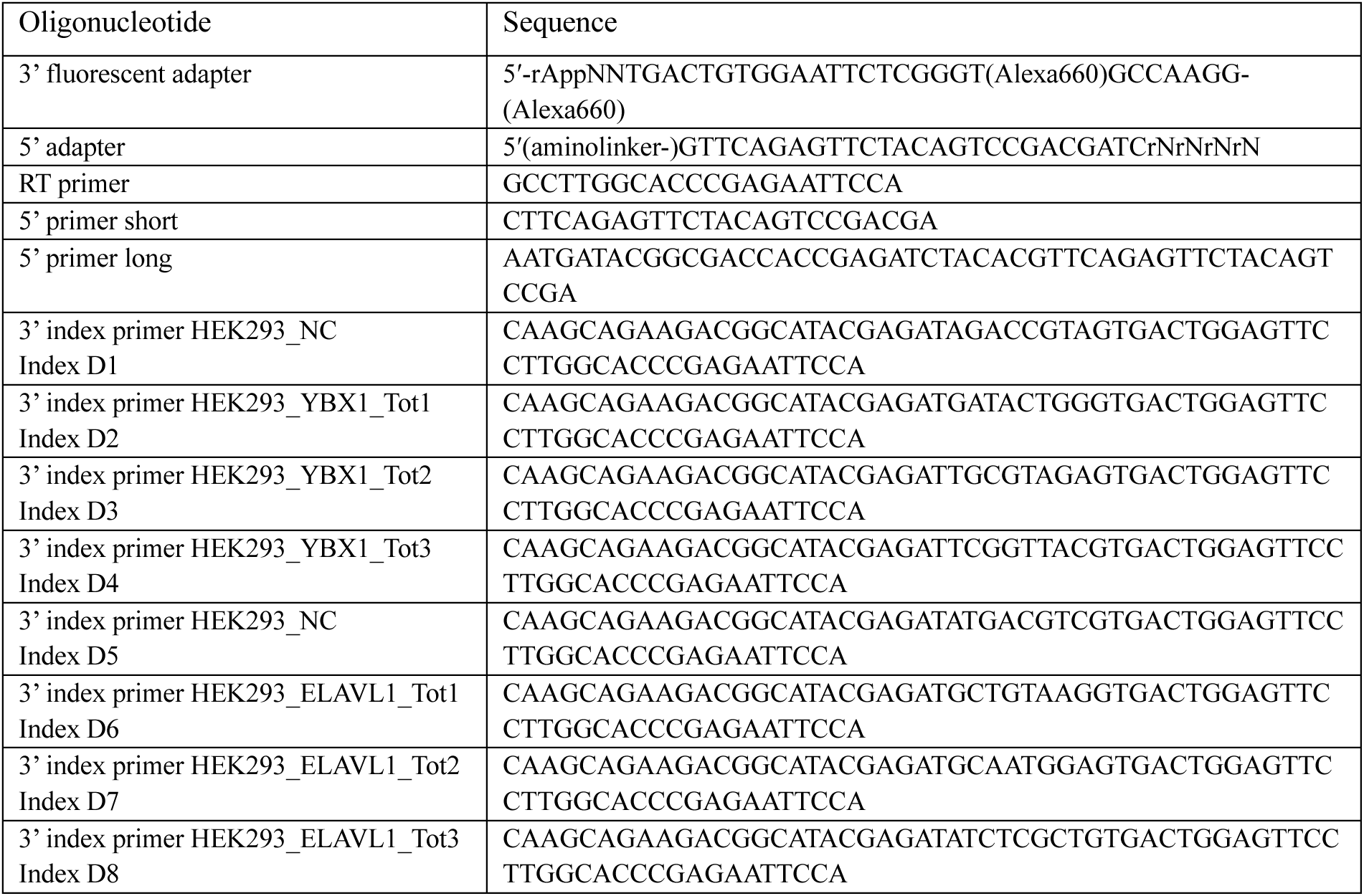
Oligonucleotides used in the study.

fPAR-CLIP data analysis was carried out using PCLIPtools (v0.7.2) (https://github.com/paulahsan/pcliptools) [62]. FASTQ files for the fPAR-CLIP experiments were assembled to hg38 version of the genome through the ‘pcliptools-align’ script of PCLIPtools. ‘pcliptools-align’ collapses the FASTQ files to FASTA files, removes duplicates, and trimmes the 3’ and 5’ adapters by using cutadapt (v4.5). The resulting FASTA files were aligned to the hg38 genome with the STAR aligner (v2.7.9a). Since adapters were removed while generating FASTA files and short read alignments were expected, an ‘end-to-end’ alignment mode was set for STAR aligner. For the alignment, the maximum number of mismatches was set to 2. Additionally, ‘--outFilterMultimapNmax’ was set to 10 to limit the maximum number of multi-mappers for a read to 10. After generating a Binary Alignment Map (BAM) file, the ‘pcliptools-cluster’ script of the PCLIPtools was used for peak calling. PCLIPtools extracts non-T-to-C mismatch information from the BAM files and fits a Poisson model to estimate background mismatch rates. Regions with observed T-to-C conversions exceeding the expected background are identified as high-confidence peaks in the fPAR-CLIP data, representing potential RNA–RBP interaction sites (clusters). An ENSEMBL-generated Gene Transfer Format (GTF) file [63] was used to annotate clusters to their corresponding genomic regions, including 5′ UTRs, coding sequences (CDS), 3′ UTRs, introns, and miRNAs. A minimum read depth of three was set for cluster inclusion. Clusters were subsequently filtered based on read depth (read_depth >= 10, the number of unique T-to-C mismatch positions (loc_T2C >= 2), and then ranked by total T-to-C counts (count_T2C).

### RBProximity-CLIP procedure 1 – FLAG–AGO2 IP first, followed by Streptavidin affinity purification

HEK293 cells expressing APEX2–fusion proteins and FLAG(HA)-tagged AGO2 (FH-AGO2-HEK293) were supplemented with 4SU and doxycycline, biotinylated, crosslinked, and lysed as described above. Subsequently, 20 μl M2 FLAG beads (Sigma, M8823) were used to immunoprecipitate tagged AGO2 from 1.5 mg of whole cell protein extracts. Next, samples were processed as described above in the fPAR-CLIP section, up to the point of SDS-PAGE and the excision of the fluorescent RNP from the gel, at approximately 130 kDa. For reconstitution into solution, gel fragments were soaked in 500 µl gel elution buffer (50 mM Tris-HCl, 150 mM NaCl, 0.1 mM EDTA, pH 7.5) supplemented with protease and phosphatase inhibitor cocktail (ThermoFisher, 78445) and incubated for 16 hours using a thermomixer at 30°C with shaking at 1000 rpm. Next, the eluates were cleared by centrifugation at 21,000 *g* for 1 minute. For verification of successful reconstitution of AGO2-RNP, 2.5 % of the eluate was saved for western blot analysis.

The 500 µl solutions containing soluble 3’ ligated AGO2-RNPs, were supplemented with 500 µl of RIPA buffer and subjected to streptavidin affinity purification using 20 µl of streptavidin magnetic beads (ThermoFisher, 88816) for 1 hour at room temperature. Beads were washed as follows: 2 times with RIPA buffer, once with 1 M KCl, once with 0.1 M Na2CO3, once with 2 M urea freshly prepared in 10 mM Tris-HCl, once with RIPA buffer, and 2 times with dephosphorylation buffer (see fPAR-CLIP section above for composition). Next, AGO2-RNA footprints were 5’ ligated, on beads, by supplementing with 0.5 µl volume of 0.5 µM 5’ adapter (MultiplexDX, MDX-O-264, see **Table 2** below), 10 U of Rnl1 T4 RNA ligase, 3 U RNase inhibitor (aabiot, 037), and shaking for 1 hour at 37°C and 800 rpm using a thermomixer. At the end of the reaction, the beads were washed twice with RIPA buffer.

**Table 2.**
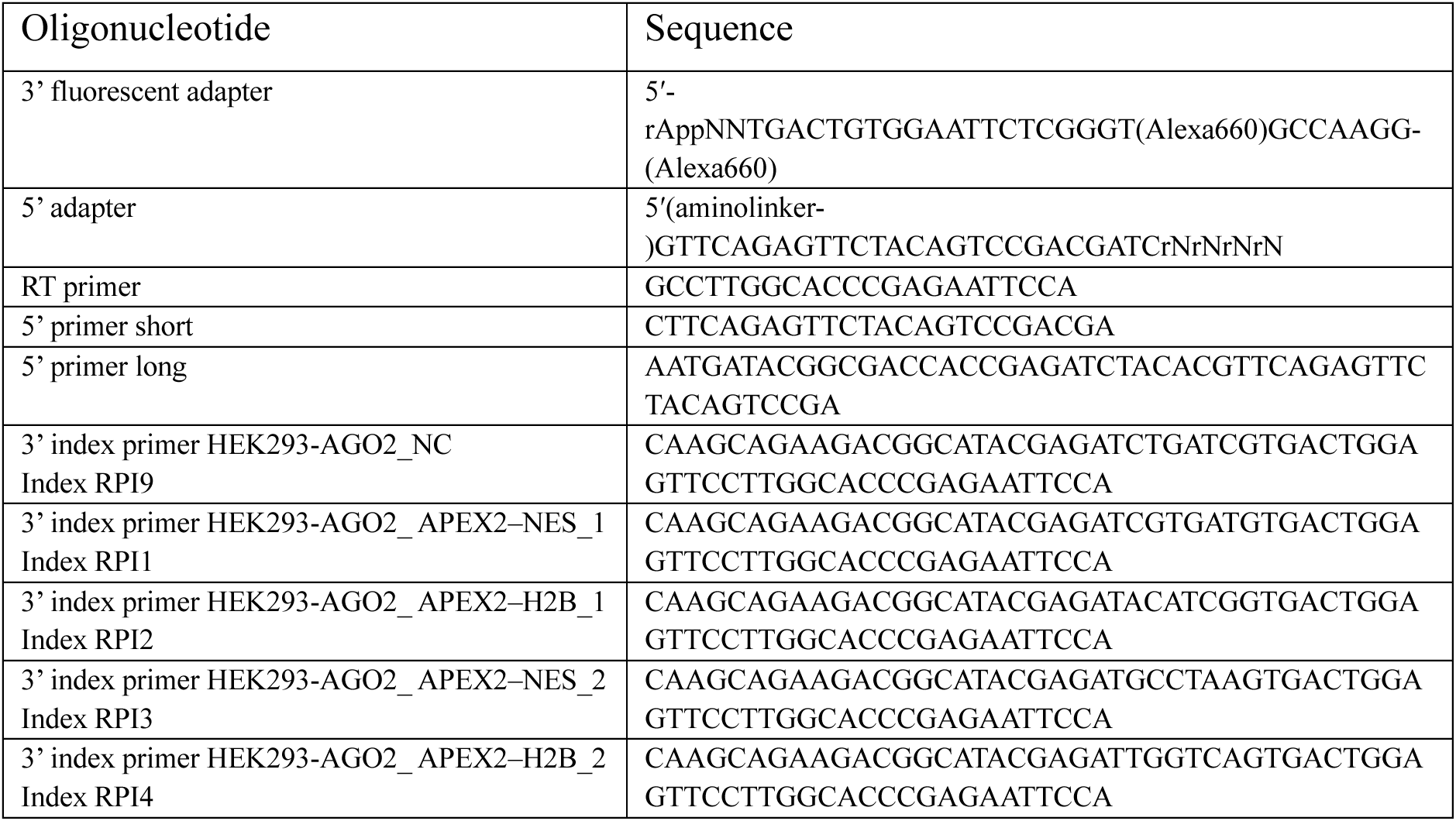
Oligonucleotides used in the study. .

To release the 3’- and 5’ ligated RNA component off the AGO2-RNPs, beads were treated with Proteinase K by stepwise additions of the enzyme, as follows: first 200 µl of 1.2 mg/ml Proteinase K in proteinase K buffer (see fPAR-CLIP section above for composition) was added and the sample was incubated for 30 minutes at 50°C with shaking at 1000 rpm. Then, 200 µl of 0.75 mg/ml Proteinase K in proteinase K buffer was added to the samples and incubated for 30 minutes in 50°C with shaking at 1000 rpm. Finally, 200 µl of 0.75 mg/ml Proteinase K in proteinase K buffer was added to the samples and incubated for 30 minutes in 50°C with shaking at 1000 rpm.

Proteinase K digestion eluates of 600 µl volume containing the released RNA footprints, were transferred to new tubes and RNA was recovered by acid phenol:chloroform extraction (ThermoFisher, AM9720). First, 1 µl GlycoBlue (ThermoFisher, AM9516), 50 µl 3 M sodium chloride, and 300 µl acidic phenol:chloroform were added to the samples. Samples were vortexed, incubated 10 minutes at room temperature, and spun down at 12,000 *g* for 15 minutes at 4°C. The aqueous (top) phase was carefully transferred to new tubes and supplemented with 200 µl water-saturated chloroform (ThermoFisher, J67241.AP). Samples were vortexed, incubated 10 minutes at room temperature, and spun down 12,000 *g* for 15 minutes at 4°C, and the aqueous (top) phase was carefully transferred to new tubes. The chloroform wash was repeated, and the RNA was precipitated by adding 1 mL of absolute ethanol (FisherBioreagents, BP2818212), vortexing, and incubating at -80°C for 1 hour. Following the incubation, samples were centrifuged at 21,000 *g* for 20 minutes at 4°C, and the RNA pellet was washed with 75 % cold ethanol. After washing, the tubes were left open for 5 minutes at room temperature to allow the ethanol to evaporate, and the RNA pellet was then dissolved in 50 µL of RNase-free water. To purify the RNA from excess salt for subsequent enzymatic reactions, the Oligo Clean and Concentrate kit (Zymo, D4061) was used. The RNA was eluted in 14 µl of RNase-free water. The eluted RNA was subjected to a reverse transcription reaction, followed by a series of PCR amplifications and size selection steps, as described in the fPAR-CLIP section above. Sequencing and data processing were carried out in a manner analogous to the fPAR-CLIP procedure.

### RBProximity-CLIP procedure 2 – biotin IP first, followed by YBX1 / ELAVL1 IP

In RBProximity-CLIP for endogenous YBX1 and ELAVL1, HEK293 cells expressing APEX2–fusion proteins were supplemented with 4SU and doxycycline, subjected to a biotinylation procedure, crosslinked, and lysed as described above for procedure 1. Here, we used 3 cell lines expressing either APEX2–NES, APEX2–H2B, or APEX2–NIK3x for proximity biotinylation of proteins in the cytoplasm, nucleus, or the nucleolus, respectively. Cell extracts corresponding to 2 mg of total protein were subjected to immunoprecipitation using 8 µg anti-biotin antibody (Abcam, ab53494) overnight. The next day, 20 µl of protein G magnetic beads (Invitrogen; 10003) were added to capture antibody-antigen complexes and incubated for 4 hours at 4°C. Then, beads were washed three times with RIPA buffer, and the bound biotinylated proteins and RNPs were eluted with 150 µl 2% (v/v) acetic acid by pipetting up and down several times. The eluates were neutralized by adding 1 M sodium hydroxide to a pH of approximately 7.5 and topped up to 1000 µl with RIPA buffer. Then the eluates were divided for YBX1 and ELAVL1 immunoprecipitation using 4 µg anti-YBX1 antibody (Abcam, 12148) and 4 µg ELAVL1 antibody (Santa Cruz, sc-5261), respectively. The antibody-antigen complexes were captured with 20 µl protein G magnetic beads for 4 hours at 4°C. After immunoprecipitation, the beads were washed three times with RIPA buffer, and the samples were eluted and processed as described in the fPAR-CLIP section above. Oligonucleotides used for YBX1, and ELAVL1 sequencing libraries preparation are listed in **Table 3** below.

**Table 3.**
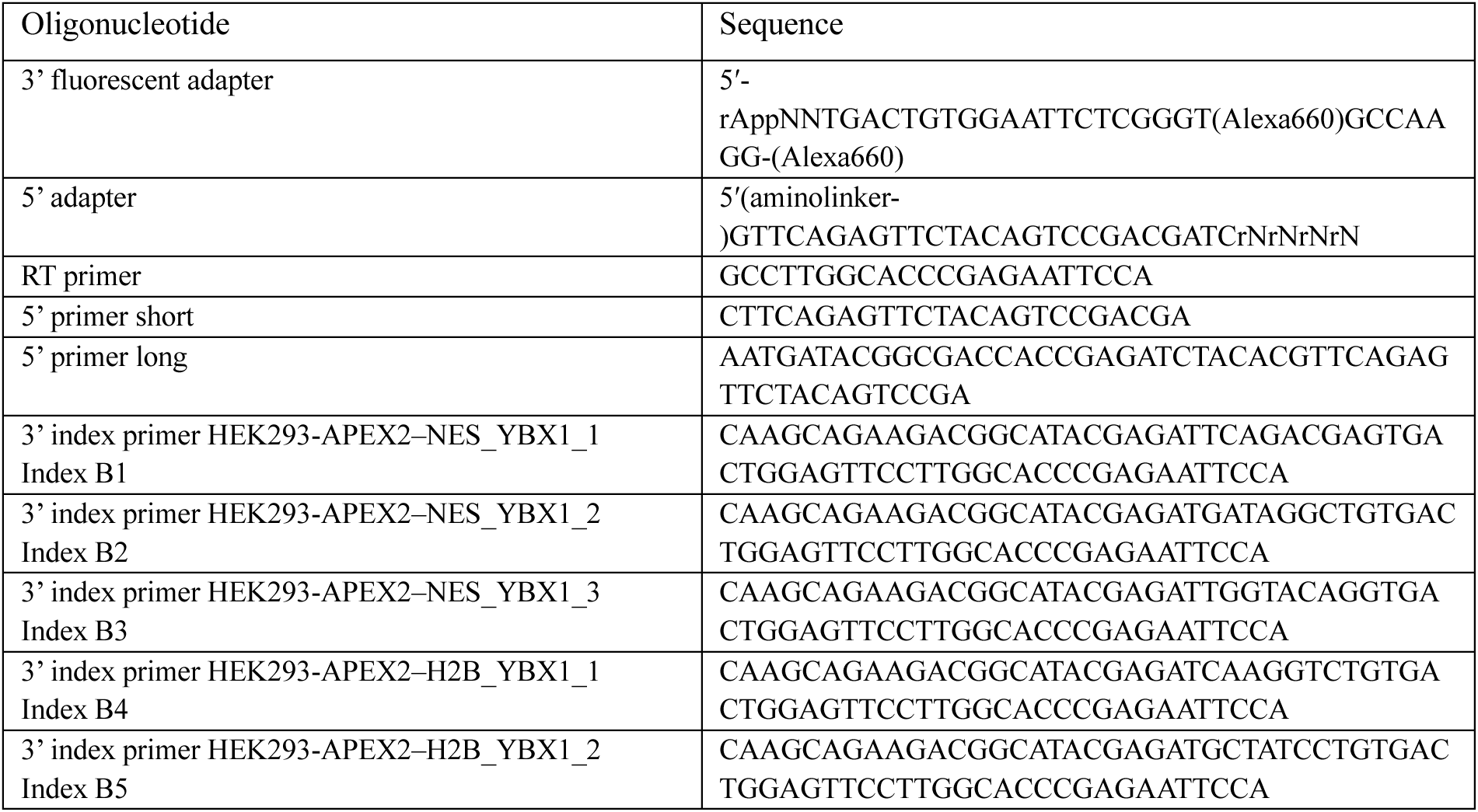
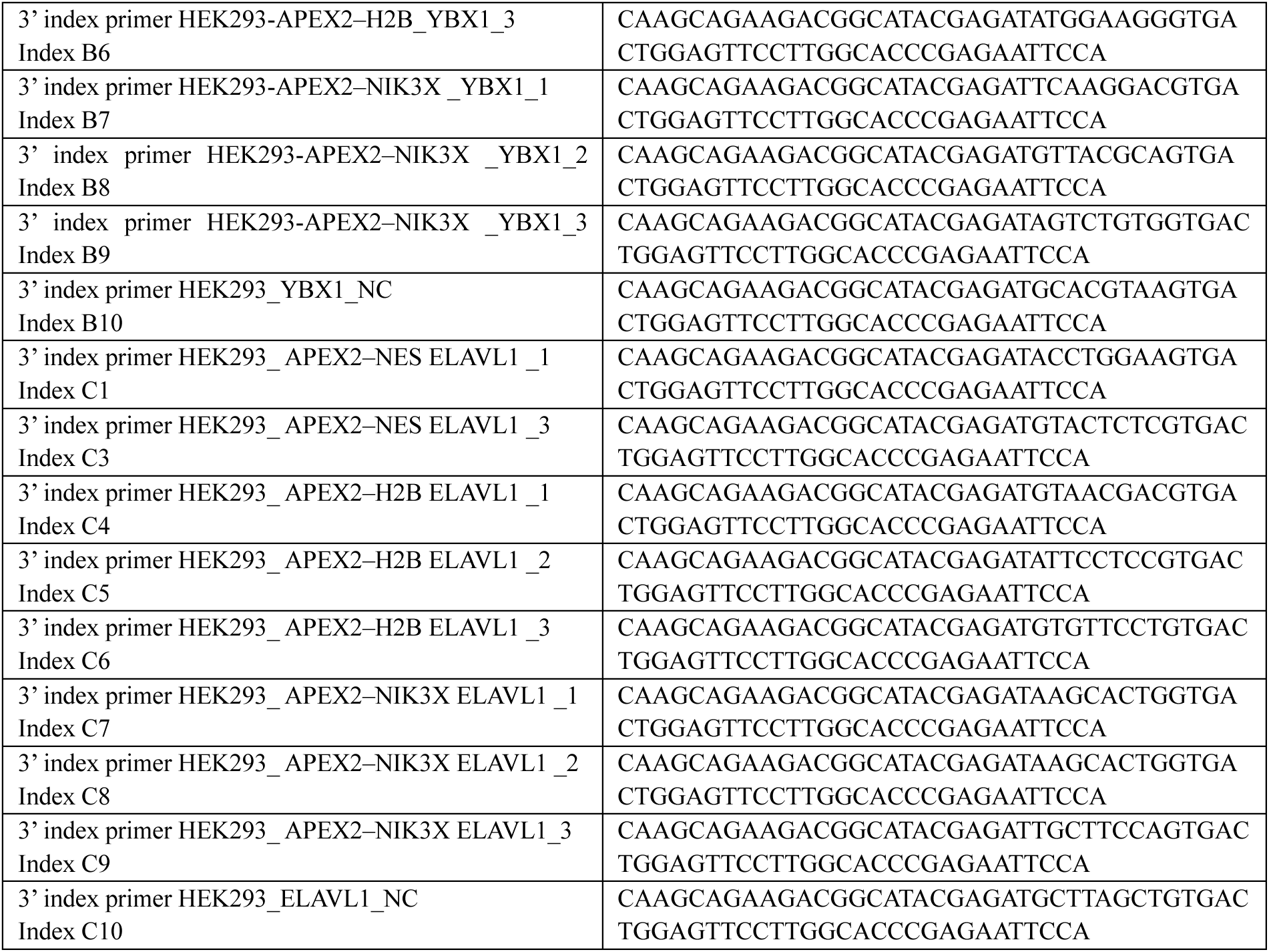
Oligonucleotides used in the study. .

### Comparison of replicates

PCLIPtools derived output files ‘clusters-stat.tsv’ were loaded into the R environment. The number of T-to-C per gene was calculated and used for pairwise comparison by means of pearson’s and/or spearman’s correlation coefficient. ‘pheatmap’ package was used to render the heatmap showing correlation. ‘pcliptools-compute-bigwig’ of PCLIPtools was used to generate ‘bigWig’ file that summarized the T-to-C mismatch information. ‘multiBigwigSummary’ and ‘plotCorrelation’ from deepTools v3.5.6 respectively were used to generate correlation matrices and plot them into heatmaps.

### Overlap and Venn diagram analysis

Intersection of YBX1 and ELAVL1 subcellular RNA interaction coordinates was performed using Bedtools intersect (v2.31.1) [64] with the -wo option, reporting all pairwise overlaps, applying a minimum overlap threshold of 1 nucleotide. The resulting output files containing feature coordinates and overlap lengths were parsed and quantified to determine shared and unique binding events across subcellular compartments. Overlap statistics were visualized using a custom Python script (Python v3.13.1) implementing matplotlib (v3.10.0) and matplotlib-venn (v1.1.2) to generate two-way Venn diagrams displaying both absolute cluster counts and relative percentages for each comparison.

### Integrative Genomics Viewer (IGV) visualization

To demonstrate compartment-specific ELAVL1 and YBX1 binding patterns, representative genomic loci were visualized using the Integrative Genomics Viewer (IGV v2.17.4) [65]. Merged fPAR-CLIP BAM files for each experimental condition were first sorted and indexed using samtools v1.22. Reads containing T-to-C conversions, indicative of productive crosslinking events, were retained using a custom Python script based on the pysam library. Normalized coverage tracks were generated with deepTools v3.5.0 [66] using the *bamCoverage* module with CPM normalization and a bin size of 1 bp to ensure consistent signal scaling across samples. The resulting CPM-normalized bigWig files were loaded into IGV alongside the GRCh38/hg38 reference genome. Visualization settings – including y-axis range, track height, and color palette – were standardized across all samples to enable direct comparison between subcellular compartments.

### 5- mer analysis

An in-house script was used to calculate the relative abundance of 5-mers in the fPAR-CLIP cluster regions, which identify overrepresented sequences in RBP-RNA interaction sites. For our analysis, 200 nt regions upstream and downstream of the clusters were used to compute background 5-mer frequencies. The script utilizes BEDtools (v2.31.1) [64] to extract FASTA sequences for both cluster regions and background regions. Afterward, the script computes a count of all possible 5-mers in both sets using Jellyfish [67]. Then the script calculates Z-score enrichment by proportion; 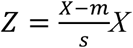 is the proportion of a given 5-mer in background regions. Additionally, m the mean and standard deviation of the respective 5-mer for the background.

## Acknowledgment

This work was funded by the Swedish Research Council (grant no. 2019-01855 to A.A.S); the Knut and Alice Wallenberg Foundation (grant no. PAR 2020/228 to A.A.S); the Swedish Society for Medical Research (grant no S19-0019 to A.A.S) and the University of Gothenburg; Intramural Research Program of the National Institutes of Health, National Institute of Arthritis and Musculoskeletal and Skin Diseases (ZIA-AR041205 to M.H.); the TAU Research and Development Fund (to D.B). We thank Dr. Stefania Dell’Orso, Faiza Naz, and Shamima Islam (NIAMS Genomics Technology Section) for high-throughput sequencing. We acknowledge the Center for Cellular Imaging at the University of Gothenburg and the National Microscopy Infrastructure, NMI (VR-RFI 2019–00217) for providing assistance in microscopy. This research was supported by the Intramural Research Program of the National Institutes of Health (NIH). The contributions of the NIH authors are considered Works of the United States Government. The findings and conclusions presented in this paper are those of the authors and do not necessarily reflect the views of the NIH or the U.S. Department of Health and Human Services.

## Author Contribution

M.H., D.B., and A.A.S., conceptualization; I.N., H.T.H., V.L., J.B.S., M.F., and G.A. methodology; I.N., A.H.P., H.T.H., M.K., V.L., J.B.S., D.G.A., M.H., D.B., and A.A.S. formal analysis; A.H.P., and M.K. data curation; I.N., D.B., and A.A.S. writing–original draft; I.N., A.H.P., H.T.H., M.K., V.L., J.B.S., M.F., G.A., D.G.A., M.H, D.B., and A.A.S. writing–review & editing; I.N., A.H.P., H.T.H., M.K., M.F., A.A.S. visualization; M.H., D.B., and A.A.S. supervision; M.H., D.B., and A.A.S. funding acquisition.

## Data availability

The raw RNAseq and fPAR-CLIP dataset described in this paper are accessible through the GEO database (https://www.ncbi.nlm.nih.gov/geo/) under accession numbers GSE311748 and GSE311747.

