## Supplemental information for "RBProximity-CLIP Enables Subcellular Mapping of RNA–Binding Protein Interactions at Nucleotide Resolution"

1 Department of Medical Biochemistry and Cell Biology, Institute of Biomedicine, University of Gothenburg, SE-40530 Gothenburg, Sweden
2 Wallenberg Centre for Molecular and Translational Medicine, University of Gothenburg, SE-40530 Gothenburg, Sweden
3 RNA Molecular Biology Laboratory, National Institute for Arthritis and Musculoskeletal and Skin Disease, Bethesda, MD 20892, USA.
4 Laboratory of Cellular RNA Biology, The Shmunis School of Biomedicine and Cancer Research, The George S. Wise Faculty of Life Sciences, Tel Aviv University, Tel Aviv, 69978, Israel
5 Nucleic Acid Therapeutics Department, Discovery Sciences, BioPharmaceuticals R&D, AstraZeneca, 43138 Gothenburg, Sweden.
6 Division of Basic Sciences, School of Medicine, University of Crete, Crete, Greece
7 The New Environmental School, Tel Aviv University, Israel.
8 Lead contact

^*^ Equal first author contribution

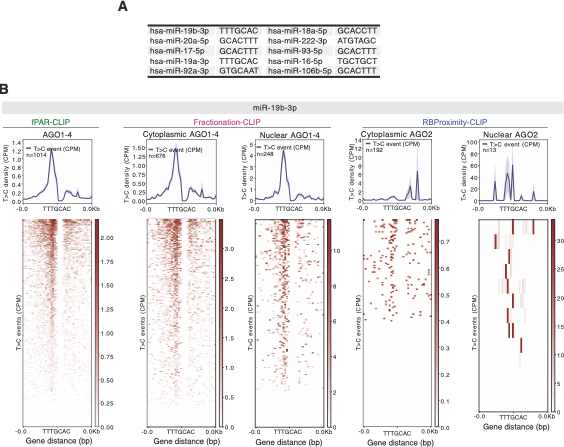

**Supplemental Figure 1 related to main Figure 2.**

**(A)** Top 10 most abundant miRNAs identified in HEK293 cells following AGO1–4 immunoprecipitation (IP). The table lists miRNA names and their corresponding seed sequences

**(B)** T-to-C conversion profile along miR-19b-3p target sites identified by AGO1–4 whole-cell fPAR-CLIP, in Fractionation-CLIP (cytoplasmic and nuclear), as well as FH-AGO2 cytoplasmic (V5-APEX2–NES) and nuclear (V5-APEX2–H2B) RBProximity-CLIP. The upper plot shows the average T-to-C conversion density (CPM) centered on the miR-19b-3p seed sequence (TTTGCAC), and the lower heatmaps display individual crosslinking events across all detected target sites (n = 1014, n =676, n = 248, n =192 and n =13).

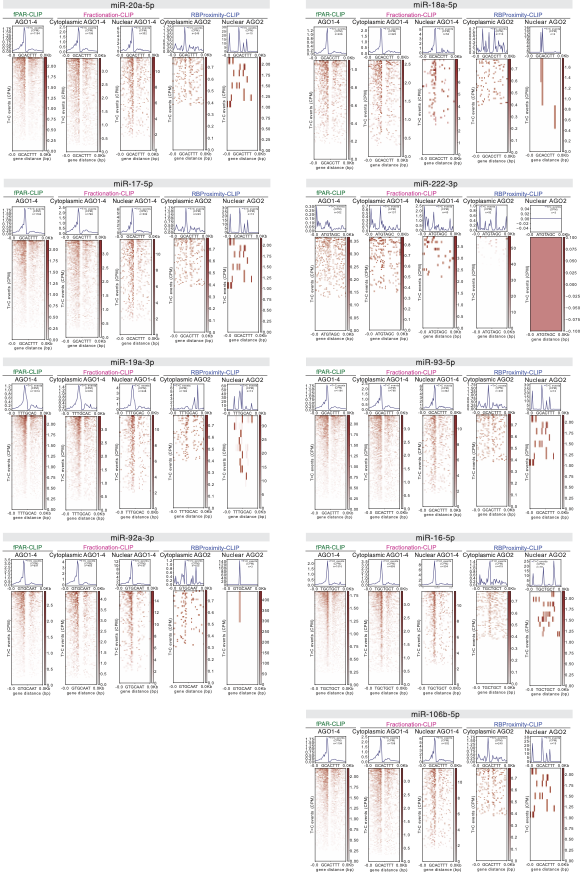

**Supplemental Figure 2 related to main Figure 2.**

Whole-cell fPAR-CLIP, cytoplasmic and nuclear Fractionation-CLIP and RBProximity-CLIP T-to-C conversion profile (CPM), cumulatively centered relative to the seed targets (top), and detailed as individual crosslinking events across all detected target sites (bottom), for the top expressed miRNAs depicted in Suppl. Fig. 1A.

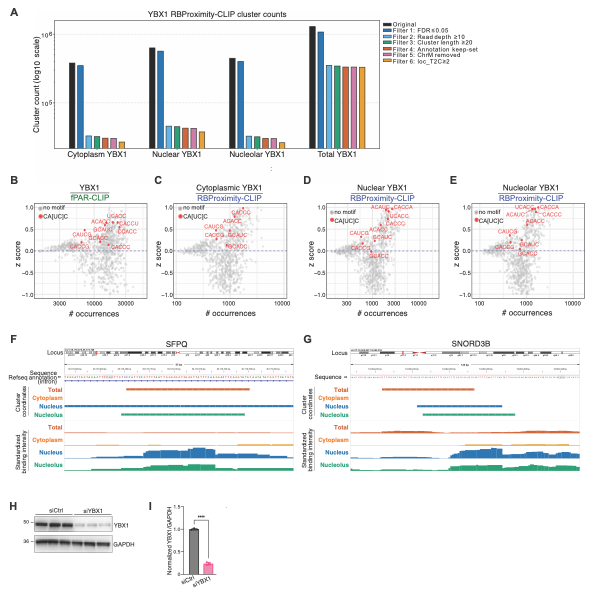

**Supplemental Figure 3 related to main Figure 4.**

**(A)** YBX1 cluster counts in response to added filtering criteria, for whole-cell fPAR-CLIP and RBProximity-CLIP samples. Filter 4 (red) retained clusters positioned along CDS, intron, UTR3, UTR5, snoRNA, miRNA, rRNA, snRNA, or lncRNA annotations.

**(B,C)** YBX1 and GAPDH Western blot analysis (**B**), quantified in (**C**), of control and YBX1 depleted HEK293 cell extracts.

**(D)** 5-mer frequency and enrichment z score (relative to a generated control) among YBX1 clusters detected by whole-cell fPAR-CLIP, cytoplasmic (V5-APEX2–NES), nuclear (V5-APEX2–H2B) and nucleolar (GFP-APEX2–NIK3x) RBProximity-CLIP. Motifs containing the CA[UC]C consensus sequence are labelled (orange).

**(E)** Genome viewer representation of standardized crosslinked reads coverage (bottom) and cluster coordinates (top) demonstrating YBX1 whole-cell fPAR-CLIP, cytoplasmic (V5-APEX2–NES), nuclear (V5-APEX2–H2B) and nucleolar (GFP-APEX2–NIK3x) RBProximity-CLIP binding, at an intronic SFPQ site and along SNORD3B.

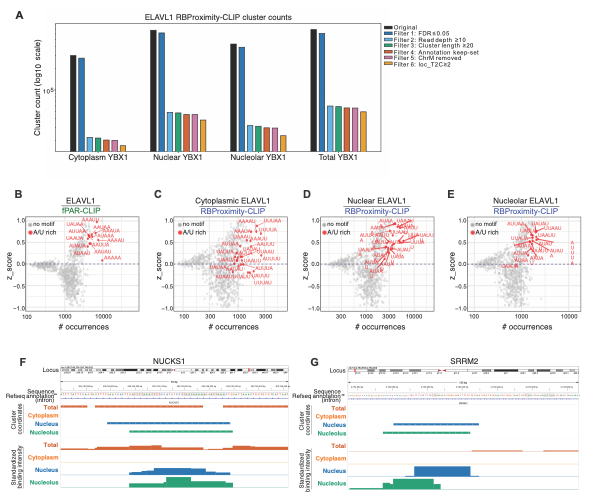

**Supplemental Figure 4 related to main Figure 5.**

**(A)** ELAVL1 cluster counts in response to added filtering criteria, for whole-cell fPAR-CLIP and RBProximity-CLIP samples. Filter 4 (red) retained clusters positioned along CDS, intron, UTR3, UTR5, snoRNA, miRNA, rRNA, snRNA, or lncRNA annotations.

**(B)** 5-mer frequency and enrichment z score (relative to a generated control) among ELAVL1 clusters detected by whole-cell fPAR-CLIP, cytoplasmic (V5-APEX2–NES), nuclear (V5-APEX2–H2B) and nucleolar (GFP-APEX2–NIK3x) RBProximity-CLIP. Motifs containing the AU-rich consensus sequence are labelled (orange).

**(C)** Genome viewer representation of standardized crosslinked reads coverage (bottom) and cluster coordinates (top) demonstrating ELAVL1 whole-cell fPAR-CLIP, cytoplasmic (V5-APEX2–NES), nuclear (V5-APEX2–H2B) and nucleolar (GFP-APEX2–NIK3x) RBProximity-CLIP binding, at intronic SRRM2 and NUCKS1 sites.
